# Multiple loci linked to inversions are associated with eye size variation in species of the *Drosophila virilis* phylad

**DOI:** 10.1101/2020.03.24.005413

**Authors:** Micael Reis, Gordon Wiegleb, Julien Claude, Rodrigo Lata, Britta Horchler, Ngoc-Thuy Ha, Christian Reimer, Cristina P. Vieira, Jorge Vieira, Nico Posnien

## Abstract

The size and shape of organs is tightly controlled to achieve optimal function. Natural morphological variations often represent functional adaptations to an ever-changing environment. For instance, variation in head morphology is pervasive in insects and the underlying molecular basis is starting to be revealed in the *Drosophila* genus for species of the *melanogaster* group. However, it remains unclear whether similar diversifications are governed by similar or different molecular mechanisms over longer timescales. To address this issue, we used species of the *virilis* phylad because they have been diverging from *D. melanogaster* for at least 40 million years. Our comprehensive morphological survey revealed remarkable differences in eye size and head shape among these species with *D. novamexicana* having the smallest eyes and southern *D. americana* populations having the largest eyes. We show that the genetic architecture underlying eye size variation is complex with multiple associated genetic variants located on most chromosomes. Our genome wide association study (GWAS) strongly suggests that some of the putative causative variants are associated with the presence of inversions. Indeed, northern populations of *D. americana* share derived inversions with *D. novamexicana* and they show smaller eyes compared to southern ones. Intriguingly, we observed a significant enrichment of genes involved in eye development on the *4*^*th*^ chromosome after intersecting chromosomal regions associated with phenotypic differences with those showing high differentiation among *D. americana* populations. We propose that variants associated with chromosomal inversions contribute to both intra- and inter-specific variation in eye size among species of the *virilis* phylad.

## Introduction

One of the most important goals of biological research is to understand the mechanisms underlying morphological diversification. The molecular basis of simple morphological traits, such as pelvic reduction in sticklebacks (Shapiro et al. 2004), presence or absence of trichomes in *Drosophila* (Sucena and Stern 2000), and pigmentation variation in flies (Wittkopp et al. 2003; Wittkopp et al. 2009) and mice (Hoekstra 2006), have been determined and are usually caused by a small number of large effect loci. However, the molecular basis of variation in complex traits remains largely elusive.

The insect head represents a great model to study complex trait evolution, since it harbors major sensory organs, which facilitate fundamental processes like feeding and reproduction. Natural variation in insect head size and shape is pervasive in insects and it is often driven by a functional trade-off between visual and olfactory sensory investment (Balkenius et al. 2006; Stieb et al. 2011; Montgomery and Ott 2015; Keesey et al. 2019; Ramaekers et al. 2019; Sheehan et al. 2019; Özer and Carle 2020), suggesting that it is likely caused by functional adaptations to an ever-changing environment. Externally, this trade-off is often observed by extensive head shape variation if compound eye size increases at the expense of the cuticle between the eyes (i.e. interstitial head cuticle) (Norry et al. 2000; Posnien et al. 2012; Keesey et al. 2019; Gaspar et al. 2020). The compound eyes are the most noticeable sensory structures in the insect head and differences in eye size have been reported between species, as well as between populations of the same species across the *Drosophila* genus (Norry et al. 2000; Hämmerle and Ferrús 2003; Posnien et al. 2012; Arif et al. 2013; Keesey et al. 2019; Ramaekers et al. 2019; Gaspar et al. 2020). Interestingly, eye size can vary due to variation in facet size or due to changes in ommatidia number (Posnien et al. 2012; Arif et al. 2013; Hilbrant et al. 2014; Gaspar et al. 2020), suggesting that different functional needs influence final eye size.

Quantitative genetics approaches have revealed multiple loci associated with variation in eye size between *D. simulans* and *D. mauritiana* supporting the complex genetic architecture of this trait (Arif et al. 2013). Similar observations were made for intra-specific variation in *D. melanogaster* (Norry and Gomez 2017; Ramaekers et al. 2019) and *D. simulans* (Gaspar et al. 2020). However, Ramaekers et al. (2019) have shown that a single mutation affecting the regulation of the *eyeless/Pax6* gene can explain up to 50% of variation in eye size between two *D. melanogaster* strains. Although, the genetic architecture underlying eye size variation is starting to be revealed for species of the *melanogaster* group, it remains unclear whether similar independent morphological diversifications identified in *Drosophila* (Norry et al. 2000; Keesey et al. 2019) share the same molecular basis over longer timescales.

Chromosomal inversions are an interesting genetic variant because suppression of recombination is thought to maintain linkage of favorable alleles which are protected from immigrant alleles carrying variants which decrease fitness (Kirkpatrick and Barton 2006; Kirkpatrick 2010). Therefore, chromosomal inversions can act as super genes influencing a myriad of phenotypes that can have a large adaptive value. The impact of chromosomal inversions on many life-history and physiological traits is well established and is often associated with local adaptation (Huang et al. 2014; Durmaz et al. 2018; Fuller et al. 2019; Kapun and Flatt 2019). Additionally, chromosomal inversions are associated with differences in rather simple morphological traits. For instance, natural variation in chromosomal inversions affect wing, thorax and head phenotypes in *D. buzzatii* (Norry et al. 1995; Fernández Iriarte et al. 2003) and wing size and shape in *D. mediopunctata* (Hatadani and Klaczko 2008) and in *D. melanogaster* (Rako et al. 2006). Chromosomal inversions are commonly associated with population structure and hinder the distinction between correlated and causative variants (Wellenreuther and Bernatchez 2018). Therefore, the impact of inversions on the diversity of complex morphological traits remains largely elusive.

Species of the *virilis* phylad of *Drosophila* are diverging from *D. melanogaster* for at least 40 million years (Morales-Hojas and Vieira 2012; Russo et al. 2013) and they have been extensively used in comparative genomics studies of important ecological traits, such as body color (Wittkopp et al. 2009; Wittkopp et al. 2011), cold resistance (Reis et al. 2011), life span (Fonseca et al. 2013) and developmental time (Reis et al. 2014). *D. virilis* is a cosmopolitan species of Asian origin while *D. americana* and *D. novamexicana* are endemic to the USA (Throckmorton 1982) and constitute the *americana* complex. *D. americana* shows a wide geographical distribution along the eastern part of the USA while *D. novamexicana* has a smaller distribution in the south-central part of the USA (Patterson and Stone 1949). Several chromosomal inversions segregate in *D. americana* populations showing latitudinal and longitudinal gradients (Hsu 1952; Throckmorton 1982). Some of these inversions create highly differentiated genomic regions between northern and southern *D. americana* populations and they are shared between northern *D. americana* populations and *D. novamexicana* (Reis et al. 2018). Since species of the *virilis* phylad, and in particular the *americana* complex, have multiple well characterized chromosomal inversions and extensive phenotypic variability, these species are a prime model to link variation in phenotypes to the presence of chromosomal inversions and simultaneously understand whether natural variation in organ morphology is due to the same molecular basis in divergent *Drosophila* lineages.

In this work we provide a comprehensive morphological and genetic characterization of eye size variation among species of the *virilis* phylad. We show that eye size differences are most pronounced between *D. novamexicana* and a southern strain of *D. americana*. Applying quantitative genetics approaches we establish that eye size differences are caused by multiple genes located in multiple chromosomes. Additionally, we found an association between the presence of chromosomal inversions and eye size. A thorough integration of population genetics, GWAS and phylogenic datasets revealed a significant enrichment for eye developmental genes among genes located on the *4*^*th*^ chromosome (Muller B). We argue that some of these variants can explain both intra- and interspecific variation in eye size.

## Results

### Head shape and eye size is remarkably variable in species of the *virilis* phylad

To evaluate the extent of variation in overall head shape in the *virilis* phylad, we performed a geometric morphometrics analysis to quantify shape differences in females of two strains of *D. virilis*, *D. novamexicana*, a northern and a southern population of *D. americana*, respectively. The mean shapes were significantly different for all possible pair-wise comparisons among species/populations (Fig 1A). We found that bigger eyes were associated with reduced interstitial cuticle and this effect was more pronounced in the ventral part of the head (Fig. 1A).

**Fig. 1.**
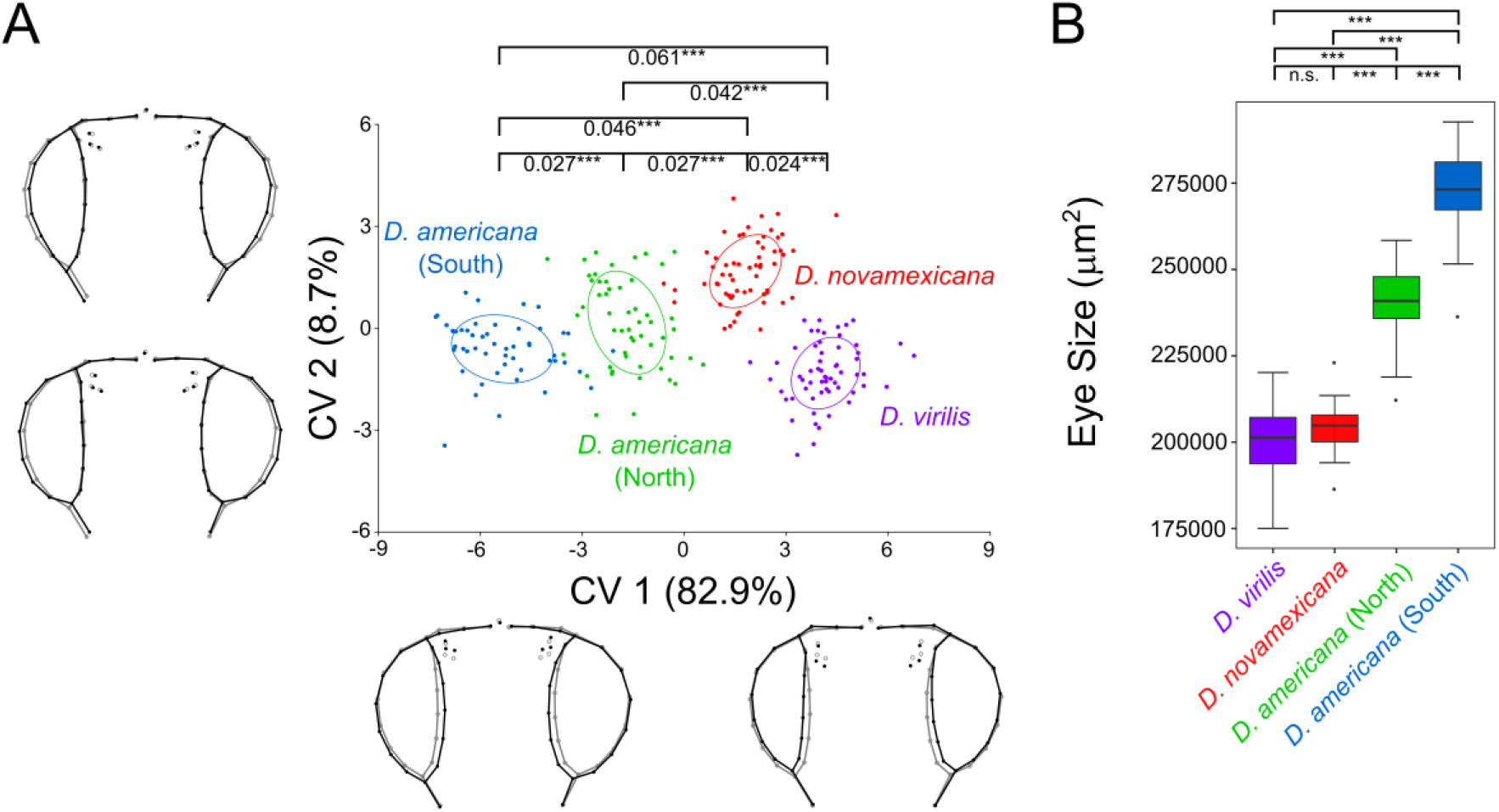
Eye size and head shape are remarkably variable among species of the *virilis* phylad. **A.** Head shape variation among species/populations (Canonical variate analysis of the procrustes coordinates obtained from the first 12 principal components (90.8% of the total variation); procrustes distances are provided with *** = p<0.0001 after a permutation test with 10,000 iterations; equal frequency ellipses are given with probability of 0.5). The wireframes depict changes in shape along the two main canonical variates (CV1 and CV2) (black - the maximum and minimum values on the axis (Mahalanobis distances); grey – mean shape for each axis). The amount of variation explained by each CV is shown in brackets. **B.** Eye size variation (after accounting to variation in body size) among species/populations (Kruskal-Wallis test, followed by post-hoc Dunn’s test and Holm correction for multiple testing: * p<0.05, *** p<0.001, n.s. p>0.05).

To confirm the observed variation in eye size, we measured eye area in females of each species/population. We observed that southern *D. americana* strains had the largest eyes while *D. virilis* and *D. novamexicana* had the smallest (Fig. 1B). Differences in eye size reached 36.2% when southern *D. americana* strains were compared to *D. virilis* and 13.7% between *D. americana* populations (File S1). Therefore, differences in head shape are accompanied by eye size (after accounting for variation in body size) variation and this association was further confirmed by the significant correlation between the former trait and CV1 (Pearson’s r = −0.925, p < 2.2e-16). Overall, these results show that eye size and head shape differ remarkably between species of the *virilis* phylad and among *D. americana* populations.

### Variation in head shape and eye size is associated with chromosomal inversions in strains of the *americana* complex

Chromosomal inversions are pervasive in the *virilis* phylad (Hsu 1952; Throckmorton 1982). Therefore, the observed variation in eye size and head shape provides an excellent model to test whether inversions are associated with differences in complex morphological traits. We developed new molecular markers for each chromosomal inversion and genotyped all analyzed strains (see Material and Methods). Our results were largely compatible with previous observations (File S2, (Hsu 1952; Throckmorton 1982)). Inversions *Xa* (Muller A) and *2a* (Muller B) were exclusive of *D. virilis*, while inversion *Xb* (Muller A) was present in all *D. novamexicana* and *D. americana* strains. Inversions *2b* (Muller E) and *5b* (Muller C) were exclusive of *D. novamexicana*, while inversion *5a* (Muller C) was exclusive of *D. americana*. Inversions *Xc* (Muller A) and *4a* (Muller B) were present in *D. novamexicana* and *D. americana* (O43, O53). For inversion *5a* (Muller C) we found evidence for heterozygosity in *D. americana* (O43) (*5a*/*5*). Surprisingly, we could not find evidence for the presence of inversion *5b* (Muller C) in northern *D. americana* strains, which was previously described to be fixed in northern populations (Hsu 1952).

Since most of the inversions, except *Xa* and *2a*, are derived in the lineage leading to *D. americana* and *D. novamexicana* (Throckmorton 1982; Reis et al. 2018), we excluded *D. virilis*, to address the impact of inversions on head shape and eye size (after accounting for variation in body size) in the *americana* complex. We found significant associations between the presence of inversions and head shape variation, mostly affecting the ratio between eye size and the head cuticle (Fig. 2A-C). Accordingly, we also found significant associations between the presence of inversions and eye size among strains (*Xbc,4a* v. *Xb,4* (Muller A, B) (W = 5489, p < 2.2e-16); *2bc,5b* v. *2,5,5a* (Muller E, C) (W = 6113, p < 2.2e-16) (Fig. 2D-F). The presence of inversions *Xc,4a* (*D. novamexicana* and northern *D. americana*) and *2b,5b* (*D. novamexicana*) resulted in a 19.1% and 20.3% reduction in eye size, respectively. Inversion *5a* (*D. americana*) led to a significant increase of 26.8% (*5a* v. *5b* (W=7, p < 2.2e-16). These results suggest that at least part of the causative variants underlying variation in eye size and head shape must be located in chromosomal inversions.

**Fig. 2.**
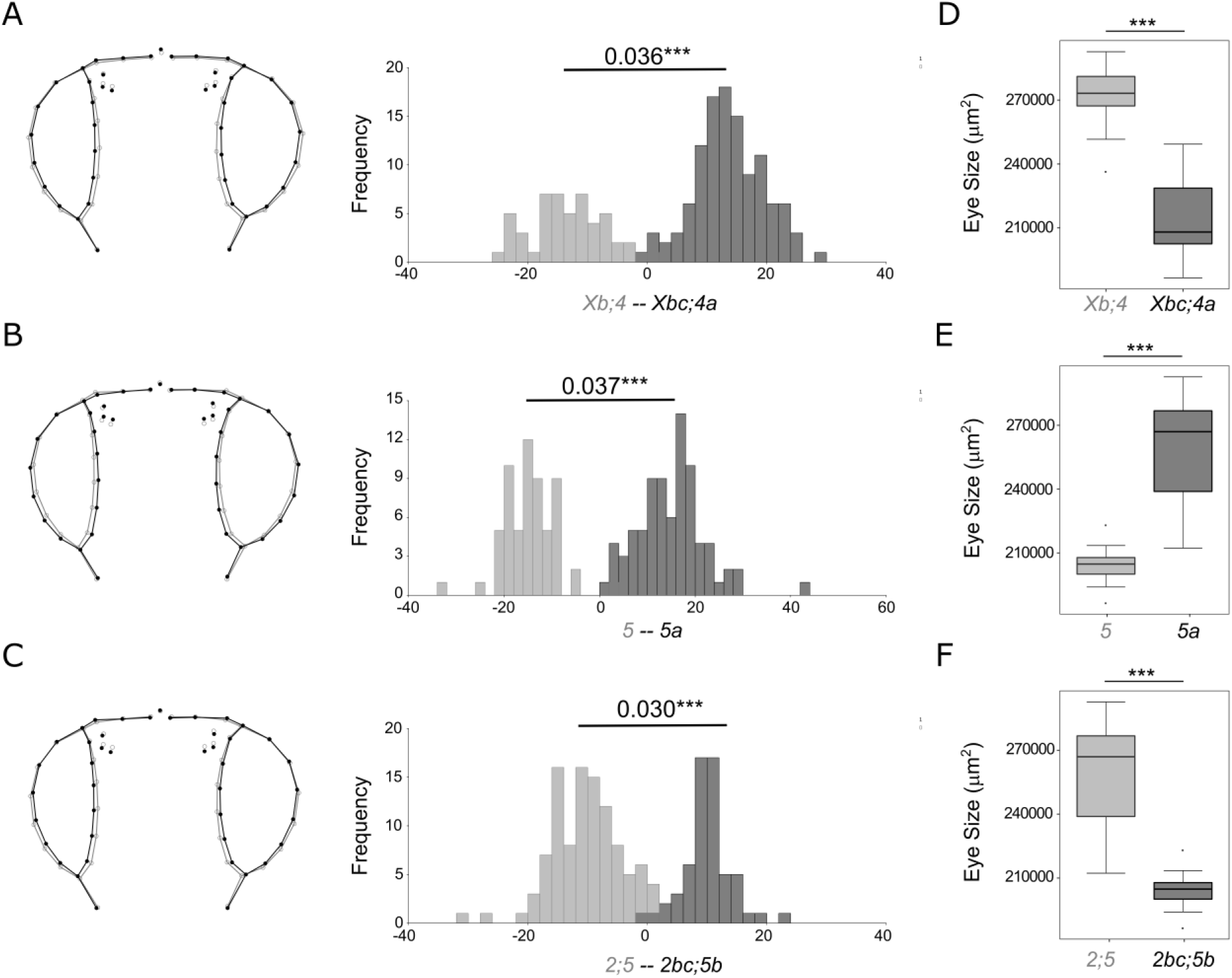
Variation in eye size and head shape is strongly associated with the presence of chromosomal inversions among species of the *americana* complex. **A-C.** Head shape variation among species/populations with (dark grey) or without (light grey) chromosomal inversions *Xc*,*4a* (D), *5a* (E), and *2bc*,*5b* (F) (Discriminant function analysis based on the Procrustes coordinates obtained from the first 12 principal components explaining 90.8% of the total variation; procrustes distances are provided with *** = p<0.0001 after a permutation test with 10,000 iterations). The wireframes depict changes in the mean shape (grey – without inversions; black – with inversions). **D-F.** Eye size variation (after accounting to variation in body size) between species/populations with (dark grey) or without (light grey) chromosomal inversions *Xc*,*4a* (A), *5a* (B), and *2bc*,*5b* (C) (Wilcoxon rank-sum test and Holm correction for multiple testing: *** p < 2.2e-6).

### Eye size is an incomplete dominant trait between *D. americana* and *D. novamexicana*

The strains showing the largest differences in eye size (after accounting for body size) were *D. americana* (SF12) and *D. novamexicana* (15010-1031.00) (File S1). Additionally, with respect to chromosomal inversions they showed the most divergent karyotypes, supporting the association between inversions and eye size. Therefore, we selected those two strains to characterize head shape and eye size variation and dominance relationship more comprehensively.

We performed a geometric morphometrics analysis to evaluate differences in head shape between both parental strains and their F1 hybrids. We found that head shapes were significantly different for all comparisons (Fig. 3A). The main differences between the species and their hybrid was explained by CV1 with the hybrid showing an intermediate head shape. CV1 captured an expansion of the eye that was accompanied by a contraction of the interstitial cuticle. This effect was more pronounced in the ventral region for *D. americana* (Fig. 3A).

**Fig. 3.**
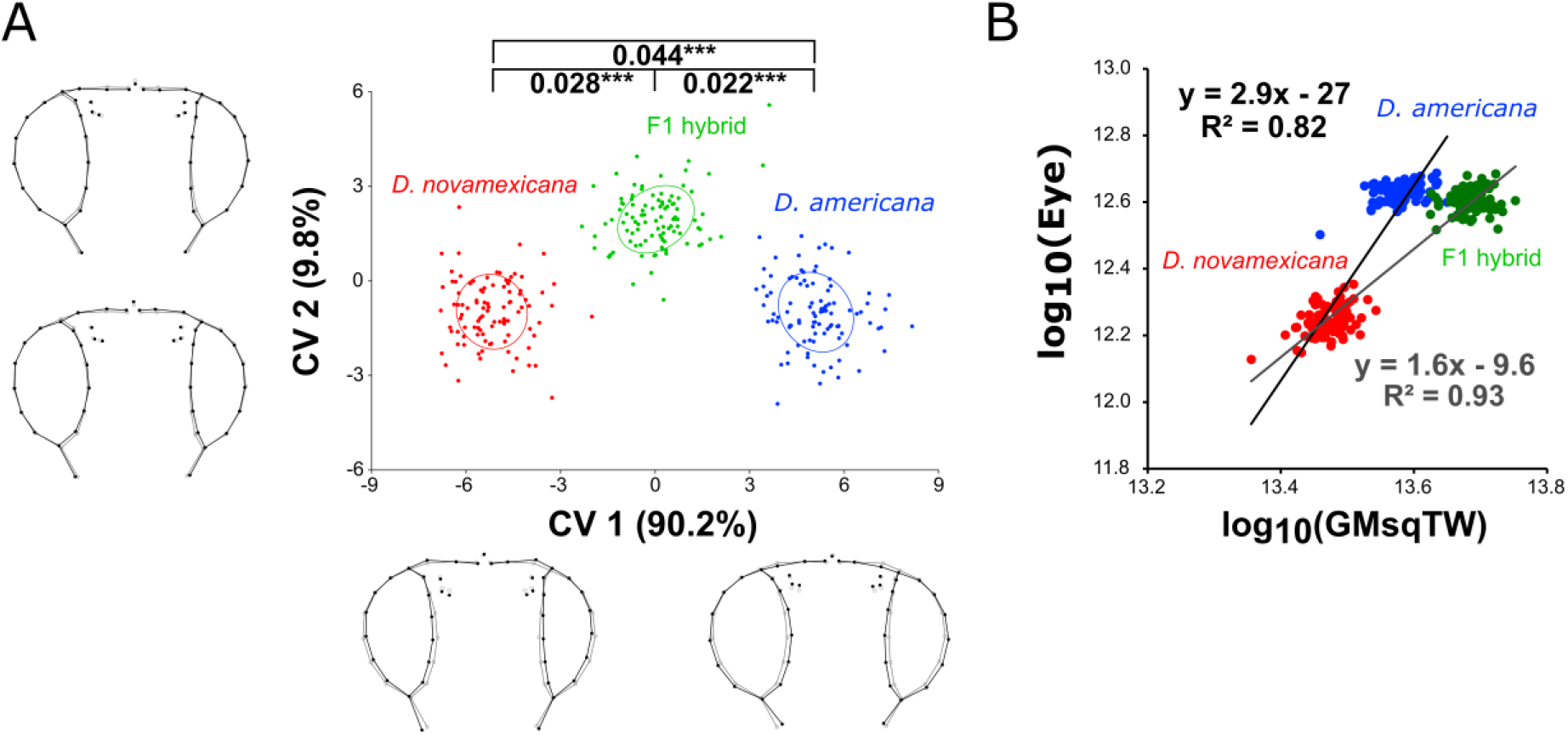
Eye size and head shape are co-dominant between *D. americana* and *D. novamexicana*. **A.** Head shape variation between parental strains (*D. novamexicana*, N=99, *D. americana*, N=100) and their inter-specific hybrids (N=100). Canonical variate analysis was applied to the procrustes distances obtained from the first nine principal components (90.5% of the total variation); procrustes distances are provided with *** = p<0.0001 after a permutation test with 10,000 iterations; equal frequency ellipses are given with probability of 0.5. The wireframes depict changes in shape along the two main canonical variates (CV1 and CV2) (black - the maximum and minimum values on the axis; grey – mean shape for each axis). The amount of variation explained by each CV is shown between brackets. **B.** Scaling relationships between females of *D. novamexicana* (15010-1031.00) (N=99) and *D. americana* (SF12) (N=98), as well as between females of *D. novamexicana* and F1 inter-specific hybrids (progeny of crosses between *D. novamexicana* males and *D. americana* females; N=98) for eye area relatively to the GMsqTW. The slopes of the equations represent the allometric coefficients between *D. novamexicana* and F1 hybrids (grey) as well as between *D. novamexicana* and *D. americana* (black).

To evaluate the dominance relationships for eye size, we compared eye areas of F1 hybrids to the parental strains and found for all comparisons statistically significant differences (Kruskal-Wallis test followed by Dunn’s post-hoc test, p < 0.001; File S1), with *D. americana* females having 46.4% larger eyes than *D. novamexicana* females. The eyes of hybrids were slightly, but significantly smaller (2.4%) than *D. americana* female eyes (File S1). Since the F1 hybrids showed almost the same size as *D. americana*, apparently the larger eyes are dominant over smaller eyes. Ommatidia counting revealed that the eye size differences were exclusively caused by variation in ommatidia number (Fig. S1).

To test whether body size influenced the observed eye size differences, we measured wing areas as well as tibiae lengths in both parental strains and in hybrids. All comparisons between the two strains and their inter-specific hybrids were statistically significant (Kruskal-Wallis test followed by Dunn’s post-hoc test, p < 0.001; File S1). Interestingly, while eye area was apparently dominant, the other organs were larger in F1 hybrids (Fig. 3B; File S1) leading to significantly larger allometric coefficients between *D. novamexicana* and *D. americana* when compared to *D. novamexicana* and F1 hybrids (B=1.30, p < 0.001; Fig. 3B). This result suggests that part of the difference in eye size between the F1 hybrids and *D. novamexicana* may be caused by pronounced changes in total body size. In summary, our results show that eye size is an incomplete dominant trait between *D. novamexicana* and *D. americana* that is largely affected by overall body size.

### Normalized eye size and head shape is affected by variation on multiple chromosomes

To reveal genetic variants associated with head shape and eye size differences between *D. americana* and *D. novamexicana*, we performed a backcross study (see Materials and Methods for details). Most chromosomes were associated with the size of multiple adult organs simultaneously with a pronounced effect of the *5*^*th*^ chromosome (Muller C) (Fig. S2), suggesting that variants in general factors affecting overall body size are segregating in these crosses. Therefore, we evaluated the effect of individual chromosomes on the non-allometric component of shape (Fig 4A-D; for the effect of each chromosome on the allometric shape see Fig. S3). We found significant associations between every chromosome and head shape variation, with smaller effects of the *X* or *5th* chromosomes (Muller A or C) compared to the *2nd*, *3rd*, and *4th* chromosomes (Muller E, D, and B) (Fig. 4A-D). The differences in mean shape caused by the *D. americana* fused *2*^*nd*^ and *3*^*rd*^ chromosomes (Muller E and D) and the *4*^*th*^ chromosome (Muller B) were compatible with a trade-off between the eyes and the interstitial cuticle (Fig. 4B, D). This effect was only observed in the ventral region of the head for *2*^*nd*^ and *3*^*rd*^ chromosomes (Muller E and D) (Fig. 4D) and the presence of the *4*^*th*^ chromosome (Muller B) additionally caused an expansion of the eye area in the lateral part of the head (Fig 4B). Hence, genes located on all chromosomes contribute to head shape variation between *D. americana* and *D. novamexicana*.

**Fig. 4.**
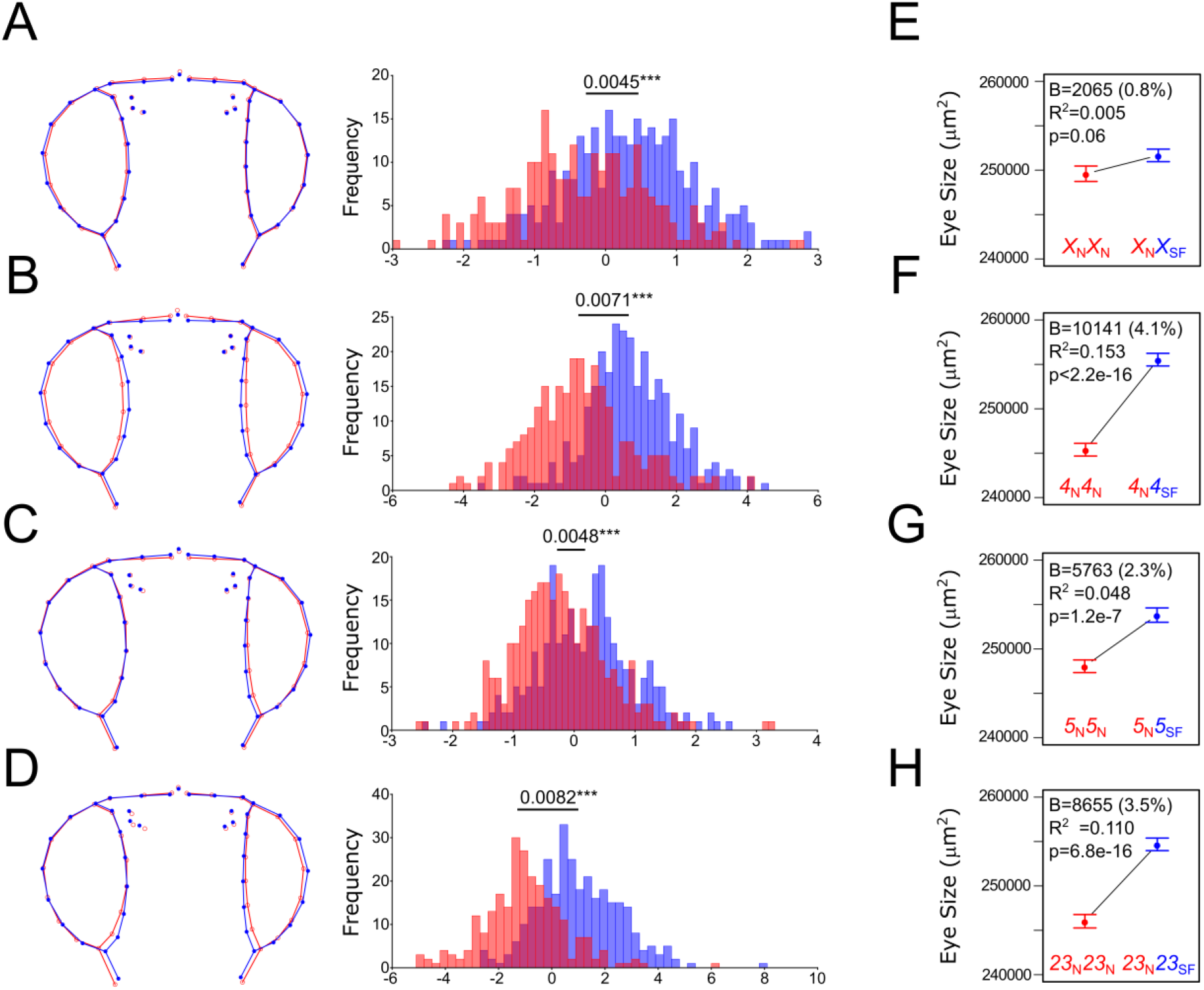
Variation in normalized eye size and head non-allometric shape is mainly explained by the *2*^*nd*^ and *3^rd^* fused chromosomes (Muller E and D) as well as by the *4^th^* chromosome (Muller B). **A-D.** Variation in mean head shape among the female progeny of the backcross between F1 hybrid females and *D. novamexicana* males (Discriminant function analysis of the procrustes coordinates obtained from the first 19 principal components (90.8% of the total variation); procrustes distances are provided with *** = p<0.0001 after a permutation test with 10,000 iterations). The wireframes depict changes in the mean shape multiplied by a factor of 5 along the axis of Mahalanobis distances (homozygous *D. novamexicana* (red) or heterozygous *D. novamexicana*/*D. americana* (blue)). **E-H** Distributions of normalized eye size (plots of means ± SEM) for females, progeny of the backcross, which were homozygous for a given *D. novamexicana* chromosome (red) or heterozygous *D. novamexicana*/*D. americana* (blue) for the respective chromosome. Information about the magnitude of change in eye size, the significance values as well as the percentage of variation explained obtained using linear regression models is shown inside the graphs.

Next, we assessed the effect of each chromosome on eye size. Since variants affecting overall body size segregated in our crosses, we determined which chromosomes affect eye size exclusively. To this end, we adopted a very conservative approach for normalization to account for variation in overall body size (see Materials and Methods for details). After accounting for differences in total body size, we observed that the *X* chromosome (Muller A) had no significant effect, while all other chromosomes showed a strong association with the normalized eye size (Fig. 4E-H). The main effects were caused by the presence of the *4*^*th*^ chromosome (Muller B) (Fig. 4F) and the fused *2*^*nd*^ and *3*^*rd*^ chromosomes (Muller E and D) (Fig. 4H), which explained 15.3% and 11.0% of the variation and resulted in an increase of 4.1% and 3.5% in normalized eye size, respectively. We also found a slight contribution of the *5*^*th*^ chromosome (Muller C) (Fig 4G) which explained 4.8% of the variation and led to an increase of 2.3% in normalized eye size.

We did not find evidence for epistasis between the chromosomes showing significant associations with normalized eye size (NormE ~ Ch23 x Ch4; NormE ~ Ch23 x Ch 5; NormE ~ Ch4 x Ch5; and NormE ~ Ch23 x Ch4 x Ch5, p > 0.05 for all interactions), suggesting that the contribution of the different chromosomes was additive. This result is further supported by the observation that the presence of the fused *2*^*nd*^ and *3*^*rd*^ chromosomes (Muller E and D), the *4*^*th*^ chromosome (Muller B) or the *5*^*th*^ chromosome (Muller C) by themselves contributed very little to an increase in normalized eye size (Fig. S4). Indeed, the genotypic class showing the highest values of normalized eye size was the one heterozygous for all chromosomes except the *X* chromosome (Muller A) (11.8% bigger than the class having *D. novamexicana* chromosomes only, File S1). We conclude that genes located on the *D. americana 2^nd^*, *3^rd^*, *4*^*th*^ and *5*^*th*^ chromosomes (Muller E, D, B, and C) when present simultaneously on a *D. novamexicana* background contribute additively to the highest increase in normalized eye size.

To increase the mapping resolution and to reveal SNPs associated with normalized eye size we performed a Genome-Wide Association Study (GWAS) using pools of individuals after 17 generations of recombination between hybrids (see Materials and Methods for details). The results obtained were highly compatible with our backcross study. We found clear regions with major differentiation between extreme quartiles on the *2*^*nd*^ and *3*^*rd*^ chromosomes (Muller E and D), as well as on the *4*^*th*^ chromosome (Muller B) and the *5*^*th*^ chromosome (Muller C) (Fig. S5). Additionally, we confirmed that the chromosomal inversions segregating in our crosses largely suppress recombination even after 17 generations. Further analysis of intermediate quartiles showed that the frequencies of the reference variants increased for different chromosomes between adjacent quartiles (Fig. S5B-D). Increased normalized eye size is, thus, caused by combinations of different chromosomes and it is largest when the frequencies of *D. americana* variants are highest across the genome. Overall, these results represent compelling evidence for the role of multiple genes located in different chromosomes in normalized eye size determination.

### Variants located in genes involved in eye development can explain both intra- and interspecific variation in normalized eye size

To narrow down the high number of potential variants (SNPs) obtained from our GWAS approach, we integrated phylogenetic and population genetics data. Under a simple additive model, the sum of the effects of the different chromosomes lead to the overall effect observed. In southern *D. americana* populations (e.g. SF12), this leads to bigger eyes while in *D. novamexicana* this leads to smaller eyes (Fig. 5A and Fig. 1B). The highly differentiated regions between northern and southern *D. americana* populations (Reis et al. 2018) should be at least partly shared between northern populations and *D. novamexicana*, because they share inversions *Xc* (Muller A), *4a* (Muller B) and *5b* (Muller C) (Fig. 5A, Fig. S6A, but see previous results). In contrast, chromosomes not showing differentiation (*2*^*nd*^ and *3^rd^*, Muller E and D) are shared between northern and southern populations (Fig. S6B) and when combined with *Xc*, *4a* and *5b* chromosomes resulted in an intermediate eye size in northern *D. americana* populations (Fig. 5A and Fig. 1B). However, the *2*^*nd*^ and *3*^*rd*^ chromosomes show extensive differentiation between *D. novamexicana* and *D. americana* alleles after 17 generations of recombination (Fig. S5), due to the presence of inversions *2b*, *2c* and *3a*, which are fixed in *D. novamexicana* (Hsu 1952; Throckmorton 1982) and contributed to a smaller eye size (Fig. 5A and Fig. 1B). Therefore, we raised the hypothesis that regions showing high differentiation due to the presence of inversions will contain the variants associated with differences in eye size in the *americana* complex.

**Fig. 5.**
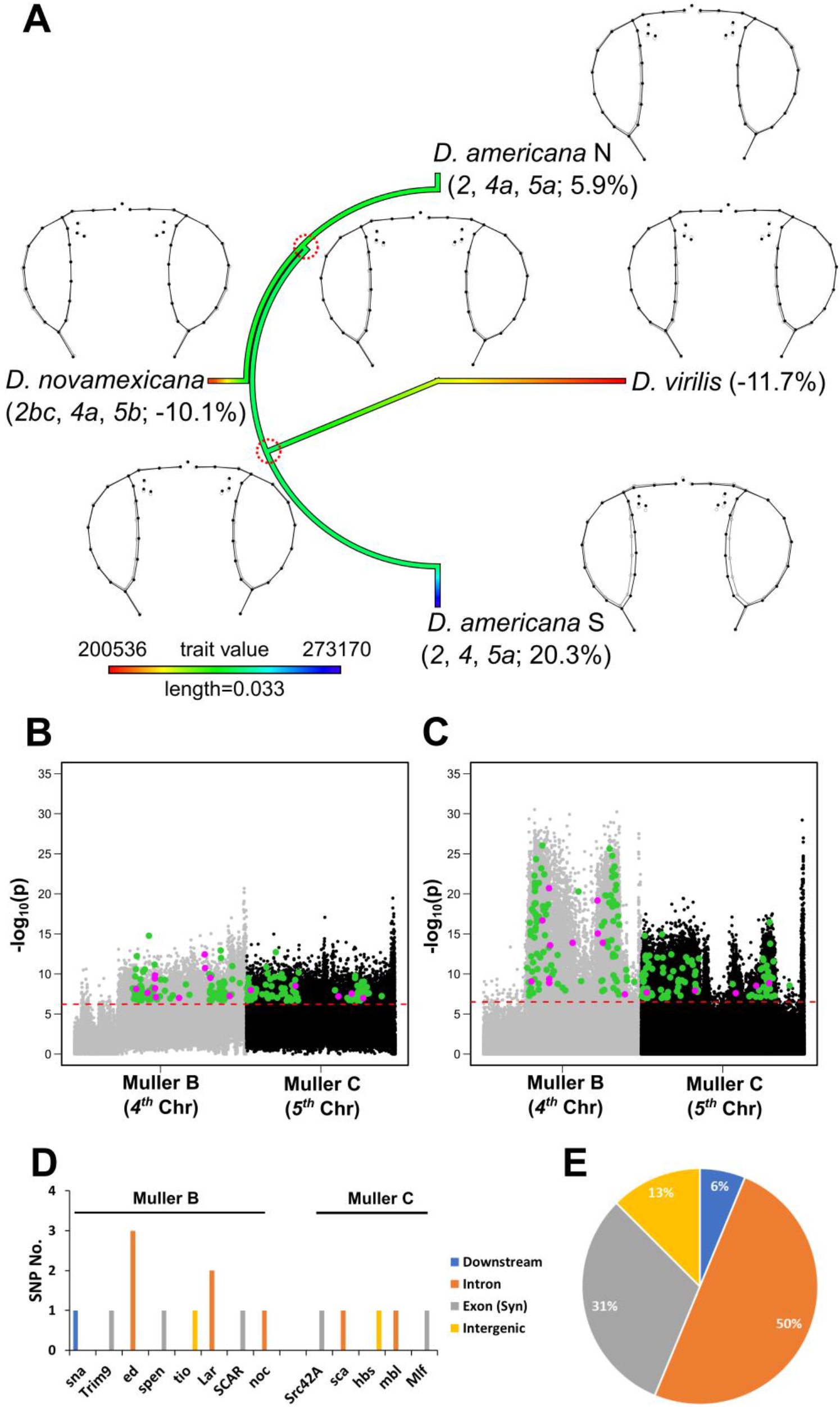
Phylogenetic and population genetics approaches revealed loci linked with inversions that can explain variation in eye size in species of the *virilis* phylad. **A.** Ancestral reconstruction of the ancestral eye size (after accounting for variation in body size) of species of the *virilis* phylad. The karyotypes, the wireframes (black – mean head shape of each species/population, grey – mean head shape of the estimated ancestral), and the percentage of variation in eye size compared to the grand mean are shown for each species/population. **B-C.** Manhattan plots of the data obtained between Q4 and Q1 (F18 pool seq) and between southern and northern *D. americana* populations, respectively for Muller B and C. The green dots depict the SNPs showing significant differences in frequencies in both data sets after Bonferroni correction; the purple dots depict the subset of significant SNPs located inside or nearby candidate genes for eye development **D.** Regions of eye development candidate genes where the SNPs depicted in purple in B-C were located. **E.** Frequency of the SNPs located in different gene regions.

Since *D. virilis* also showed smaller eye size, we reconstructed the ancestral state of size traits to understand how these phenotypes evolved in this group of species. We observed that the ancestral eye size was intermediate (Fig. 5A) and this result was supported by the ancestral reconstruction of size phenotypes of 59 *Drosophila* species reported in (Keesey et al. 2019) (Fig. S7). Given the high levels of phenotypic and nucleotide variation characteristic of *D. americana* (Fonseca et al. 2013), it is likely that the intermediate ancestral eye size represented the mean of a quantitative trait including smaller and bigger eyes. Thus, we assumed that the ancestral population of the *virilis* phylad was highly polymorphic for eye size, and both bigger and smaller eye size were selected for in specific lineages from a pool of standing genetic variation (Fig 5A). Under this hypothesis, the variants responsible for increased eye size should be fixed in the *D. americana* (SF12) reference genome and at higher frequencies in southern populations, while the alternative variants should be fixed between *D. novamexicana* and *D. virilis*. The SNPs matching these conditions and showing significant differences in frequency between extreme quartiles of the GWAS between *D. americana* and *D. novamexicana* represent prime candidates to explain eye size variation in species of the *virilis* phylad.

To obtain a list of candidate SNPs (see Fig. S8 and Materials and Methods for details), we intersected the SNPs showing significantly higher frequency of the reference variant (*D. americana* (SF12)) in Q4 compared to Q1 with those showing the reference variants at higher frequencies in southern *D. americana* populations (Reis et al. 2018). From these SNPs, we kept only those that showed the alternative variant common between *D. novamexicana* and *D. virilis*, and we annotated the SNPs to genes using the information available for *D. virilis*. We obtained a total of 6,670 SNPs within or close to 3,006 unique genes. However, only 2,627 unique genes of *D. virilis* have recognizable orthologs in *D. melanogaster* (1,254; 534; 366 and 473 on the *2*^*nd*^, *3^rd^*, *4*^*th*^ and *5*^*th*^ chromosomes (Muller E, D, B and C), respectively; File S3). Next, we identified the orthologs of these 2,627 *D. virilis* genes in *D. melanogaster* (2,701 genes due to some ambiguities, File S3) and intersected them with the 397 genes associated with eye development (see Methods). We obtained 446 SNPs located within or close to 126 unique genes (59, 22, 21 and 24 on the *2*^*nd*^, *3^rd^*, *4*^*th*^ and *5*^*th*^ chromosomes (Muller E, D, B, and C), respectively File S3). These results showed that one fifth of the total number of genes estimated to be present in *D. melanogaster* (2,701/13,767) had SNPs with significant frequency differences in the GWAS and the same variant in *D. novamexicana* and *D. virilis*, but a different variant in *D. americana* (SF12). Almost one third of the total number of candidate genes for eye development (126/397) were present among those genes. Thus, in our dataset we observed a significant over-representation of eye developmental genes (126 out of 2,701 vs. 397 out of 13,701, Chi-square statistics with Yates correction = 37.25, p<1.00e-5) that may explain variation in normalized eye size among species of the *virilis* phylad.

We also postulated that the combination of different variants shared either with *D. novamexicana* or with southern *D. americana* populations leads to an intermediate phenotype in northern *D. americana* populations (Fig. 5A). Thus, the overlap between the chromosomal regions associated with phenotypic differences with those showing high differentiation among *D. americana* populations (Reis et al. 2018) may explain both inter- and intra-specific variation in eye size. The intersection of both datasets resulted in a total of 119 SNPs in 102 genes on the *4*^*th*^ chromosome (Muller B) and 101 SNPs in 102 genes on the *5*^*th*^ chromosomes (Muller C) (File S3). After intersecting these 204 genes with the 397 candidates for eye development (see methods), we obtained 11 SNPs in 8 out of 102 genes on the *4*^*th*^ chromosome, and 5 SNPs on 5 out of 102 genes in the *5*^*th*^ (Fig 5B-C). There was a significant over-representation of genes involved in eye development on the *4*^*th*^ chromosome (8 out of 102 v. 397 out of 13,701; Chi-square statistics with Yates correction = 7.13, p = 7.6e-3), but this was not the case for the *5^th^* (5 out of 102 v. 397 out of 13,701; Chi-square statistics with Yates correction = 0.84, p = 0.36) (Fig 5 D). The majority of the identified SNPs were in non-coding regions (69%), suggesting that regulatory sequences and thus associated gene expression may be predominantly affected (Fig. 5E). As expected, most of the SNPs that can explain both intra- and interspecific variation in normalized eye size were located on the *4*^*th*^ and *5*^*th*^ chromosomes (Muller B and C). However, we also observed two SNPs in two genes on the *2*^*nd*^ chromosome (Muller E) and four SNPs in four genes on the *3*^*rd*^ chromosome (Muller D). One of the SNPs on the *3*^*rd*^ chromosome was a synonymous mutation located in one candidate gene for eye development) (File S3). In summary, we revealed a significant enrichment in genes involved in eye development on the *4*^*th*^ chromosome carrying variants strongly associated with normalized eye size variation between species of the *virilis* phylad and within *D. americana*.

## Discussion

We provide the most comprehensive morphological and molecular characterization of head shape and eye size variation in species outside the *melanogaster* group. We observed remarkable differences in these two traits among species of the *virilis* phylad. Our shape analysis revealed that increased eye size was accompanied by a contraction of the interstitial head cuticle. A similar trade-off has been observed in other *Drosophila* species (Norry et al. 2000; Posnien et al. 2012; Arif et al. 2013; Keesey et al. 2019; Ramaekers et al. 2019; Gaspar et al. 2020) and it may be associated with different investment in visual or olfactory sensory perception (Keesey et al. 2019; Ramaekers et al. 2019). Indeed, it has been shown that a northern *D. americana* strain is more “visual” because it has significantly bigger eyes compared to the antennae, while *D. virilis* is a more “olfactory” species with smaller eyes and bigger antennae (Keesey et al. 2019). Following this rational, *D. novamexicana* may be a more “olfactory” species compared to the southern *D. americana* strain studied in detail here.

While differential investment in visual or olfactory sensory information is a common phenomenon in animals (Balkenius et al. 2006; Stieb et al. 2011; Montgomery and Ott 2015; Keesey et al. 2019; Ramaekers et al. 2019; Sheehan et al. 2019; Özer and Carle 2020), it remains to be established whether the genetic underpinnings are the same or not. Since the compound eyes (i.e. vision) and the antennae (i.e. olfaction) originate from the same imaginal disc during larval development (Haynie and Bryant 1986), the pervasive variation in head shape and eye size in *Drosophila* is an excellent model to test this. Variation in eye size between *D. melanogaster* strains is highly associated with one SNP in the Cut transcription factor binding site in the *eyeless/Pax6* regulatory region (Ramaekers et al. 2019). However, neither *eyeless/pax6* nor *cut* can explain the natural variation in eye size observed among species of the *virilis* phylad, because they are located on chromosomes that are not associated with differences in eye size among these species in our study. Also, data obtained for *D. mauritiana* and *D. simulans* showed that differences in eye size are due to variation in facet size (Posnien et al. 2012), while eye size differences between *D. novamexicana* and *D. americana* were exclusively caused by differences in ommatidia number. Since facet size and ommatidia number are specified by different developmental processes (Şahin and Çelik 2013; Treisman 2013), it is likely that different developmental mechanisms underly natural variation in eye size. Indeed, the most significant QTL explaining eye size variation in *D. mauritiana* and *D. simulans* mapped to the *X* chromosome (Arif et al. 2013). In contrast, our data showed that genetic variants affecting exclusively eye size were located in all chromosomes, but not in the *X* and *6*^*th*^ chromosomes (Muller A and F). Therefore, our comparative morphological and mapping data strongly suggest an independent evolution of eye size in different lineages. This observation is supported by similar data obtained for two species of the *melanogaster* group (Gaspar et al. 2020).

The imaginal disc that gives rise to the *Drosophila* head is a modular structure that contributes cells to almost all organs of the head (Haynie and Bryant 1986). Since we observed an additive effect of *D. americana* chromosomes in a *D. novamexicana* background on eye size, it is conceivable that each chromosome or chromosomal region might influence different developmental processes and different organ anlagen within the imaginal disc. This hypothesis is supported by our observation that the *D. americana 5*^*th*^ chromosome (Muller C) had a major impact on the size of all organs in our study, while the *2*^*nd*^, *3*^*rd*^ and *4*^*th*^ chromosomes (Muller E, D and B) were associated with variation in eye size after accounting for body size. Please note that we cannot rule out the presence of variants located on the *5*^*th*^ chromosome (Muller C) affecting exclusively eye size that were masked by the conservative approach for body size correction used in this work. Our shape analysis further revealed that only the *4*^*th*^ chromosome (Muller B) was associated with variation in lateral eye regions, supporting a modular impact of different chromosomes on overall head shape variation. For species of the *melanogaster* group it has also been shown that the evolution of eye size and the size of the interstitial cuticle is uncoupled (Arif et al. 2013; Gaspar et al. 2020). In contrast, the SNP in the *eyeless/Pax6* locus associated with intra-specific variation in *D. melanogaster* influences the early subdivision of the imaginal disc into the retinal and the antennal part of the imaginal disc (Ramaekers et al. 2019). Therefore, this SNP may affect eye size and head cuticle/antennal size simultaneously. Although more comparative developmental analyses are necessary, the picture emerges that the modular nature of the imaginal disc with its different interconnected developmental programs may facilitate the independent evolution of head shape because it provides multiple targets for evolutionary changes.

In our survey we observed the most pronounced differences in eye size between *D. novamexicana* showing the smallest eyes and southern *D. americana* strains showing the largest eyes. Interestingly, compatible with studies using species of the *melanogaster* group (Norry et al. 2000; Posnien et al. 2012; Norry and Gomez 2017; Ramaekers et al. 2019; Gaspar et al. 2020), we also identified intra-specific differences among *D. americana* populations. There are multiple chromosomal inversions segregating in *D. americana* populations and some of them are shared with *D. novamexicana* (Hsu 1952; Throckmorton 1982). These inversions largely affected the patterns of differentiation along chromosomes among *D. americana* natural populations (Reis et al. 2018). Although the genomic structure caused by the presence of inversions hampers the identification of causative variants, it has been proposed that they may keep together favorable combinations of alleles (Kirkpatrick and Barton 2006; Kirkpatrick 2010). Interestingly, we observed a clear association between the presence of shared inversions and eye size. Additionally, we found a significant enrichment of genes involved in eye development among those genes containing SNPs that could explain both intra- and inter-specific differences in eye size for the *4*^*th*^ chromosome (Muller B) only. These genes represent prime candidates for future functional validation tests.

Since these inversions show latitudinal and longitudinal gradients in the *americana* complex, it is likely that they carry the targets of selection associated with local adaptation in natural populations. For instance, chromosomal inversions were found to be associated with life-history and physiological traits likely involved in adaptation (Huang et al. 2014; Durmaz et al. 2018; Kapun and Flatt 2019) as well as with morphological traits (Norry et al. 1995; Fernández Iriarte et al. 2003; Rako et al. 2006; Hatadani and Klaczko 2008). Additionally, a previous study found that the fixed variant explaining pigmentation differences between *D. novamexicana* and *D. americana* was polymorphic in *D. americana* and explained the least pronounced variation in pigmentation observed along a longitudinal transect in *D. americana* populations (Wittkopp et al 2009). Therefore, it is conceivable that inter-specific differences affecting exclusively eye size between *D. novamexicana* and *D. americana* can also explain intra-specific differences in this trait among *D. americana* natural populations. The longitudinal gradient for pigmentation in *D. americana* populations was further confirmed (Wittkopp et al. 2011), and solar radiation and the diurnal temperature range has been shown to be the best predictors of this gradient (Clusella-Trullas and Terblanche 2011). Interestingly, *D. americana* populations showing darker pigmentation were more often found in geographical regions with lower sun radiation and mean diurnal temperature ranges than lighter populations (Clusella-Trullas and Terblanche 2011). According to these geographical parameters, we observed here that flies showing bigger eye size, likely more sensitive to light than flies with smaller eyes, came from regions with low sun radiation and possibly with less light. Therefore, eye size variation in the *americana* complex may be associated with local adaptation as well.

In conclusion, natural variation in head morphology is common in *Drosophila* and has a strong genetic component. We provide for the first time a comprehensive morphological comparison of eye size and head shape between *D. novamexicana* and *D. americana* and revealed a complex underlying genetic architecture. Our data strongly suggests that the presence of inversions in these two species contributed to nucleotide diversity patterns that may affect the regulation and function of multiple genes during head and eye development and thus facilitating natural variation in this complex morphological trait.

## Materials and Methods

### Fly strains

The following isofemale fly strains were used in this work: *D. virilis* (15010-1051.47, Hangzhou, China; 15010-1051.49, Chaco, Argentina), *D. novamexicana* (15010-1031.00, Grand Junction, Colorado, USA; 15010-1031.04, Moab, Utah, USA) and *D. americana* (O43, O53, SF12 and SF15). The *D. virilis* and *D. novamexicana* strains were obtained from the Tucson stock center in 1995 and were kept in the laboratory since then. *D. americana* strains were established with single inseminated females collected from the wild in different locations of the USA (Omaha, Nebraska (O), 2008 and Saint Francisville (SF), Louisiana, 2010) (Reis et al. 2011; Fonseca et al. 2013; Reis et al. 2015). All strains were kept at 25°C under 12h/12h light/dark cycles.

### Dissection and phenotyping

To study size variation, we dissected heads, wings and tibiae of 20-30 females between 4 and 7 days after eclosion for each strain. To avoid crowding effects on adult organ size, we controlled for density by transferring 30 first instar larvae into single vials containing standard food. The heads were mounted facing upwards on a slide with sticky tape, while the three legs (one of each pair) and wings were randomly dissected from the left or right side and mounted on a slide with Hoyer’s medium. Pictures were taken using a stereomicroscope Leica M205 FA with a magnification of 50x for wings and 60x for the other structures. We also took a picture of a ruler to be able to convert the measurements from pixels to μm or μm^2^. The resulting JPG files were saved with a resolution of 2560×1920 pixels, and we used ImageJ (Schneider et al. 2012) to measure eye areas as well as tibiae lengths and wing areas (Fig. S9A, File S4). We calculated the geometric mean of squared tibiae (GMsqT) as a proxy for tibiae size. The geometric mean of squared tibiae and wing area (GMsqTW) was used to estimate overall body size. We then used the residuals of the linear regression between eye size and GMsqTW to account for differences in overall body size between the strains. Since the measurements were not normally distributed (Shapiro-Wilk, p < 0.05), we used Kruskal-Wallis test followed by Dunn’s post-hoc test with Holm correction to determine which comparisons were significantly different between strains.

### Geometric morphometrics

Frontal head images of every strain were used to generate tps files in which all images were randomized with tpsUtil (version 1.60; (Rohlf 2015)). These tps files were used to place 43 landmarks and semilandmarks (see Fig. S9B) using tpsDig2 (version 2.18; (Rohlf 2015)). A sliders file that contains information about the semilandmarks was generated with the “Make sliders file” function in tpsUtil (version 1.60; (Rohlf 2015)). Using tpsRelw (version 1.57, 64 bit; (Rohlf 2015)) the semilandmarks were slid along a curve using an option to minimize the bending energy required for a deformation of the consensus to the selected specimen (Slide method = Chord min BE) allowing up to three iterations during the superimposition process (Slide max iters = 3). The slid landmarks were treated as fixed landmarks and were superimposed using Procrustes fit as implemented in MorphoJ (version 1.06d; (Klingenberg 2011)). Since we used 2D pictures of 3D structures, after a principal component analysis (PCA) we observed that artificial pitch (up/down rotation) and yaw (left/right rotation) were partly associated with shape variation along principal component (PC) 1 and PC3 axes, respectively (Fig. S10). Additionally, we observed that part of the within-strain variation captured pitch and yaw. To dissociate and remove the error from the true components of shape, we used a two-step approach: First, we calculated the residuals of the within-strain pooled-regression between Procrustes coordinates and PC1. Second, we used the new coordinates to repeat the above-mentioned procedure to remove the effect of yaw (new PC2). To avoid inflation of the number of the variables when compared to the number of samples in the statistical analysis, we used the final Procrustes coordinates to determine and keep the number of PCs explaining about 90% of the total shape variation. The new dataset was rotated back into the original Procrustes coordinates by transposing the orthogonal matrix. Differences in mean shape among species/strains were evaluated using a T-square parametric test followed by 10,000 permutations (leave-one-out cross-validation) as implemented in the “Canonical Variate Analysis” option in MorphoJ. The error correction and wireframes generation were performed with MorphoJ while the PC removal and rotation to the original coordinates was done using a custom R script.

### Impact of chromosomal inversions on eye size

To test whether the presence of inversions was associated with differences in eye size (after accounting for body size) and head shape, we screened *D. americana*, *D. novamexicana* and *D. virilis* strains for the presence/absence of eight different inversions known to be fixed or polymorphic within the *virilis* phylad of *Drosophila* (Hsu 1952; Throckmorton 1982) and for which the breakpoint locations have been identified (*Xa*, *Xb*, *Xc* (Muller A), *2a*, *2b* (Muller E), *4a* (Muller B), *5a* and *5b* (Muller C), (Evans et al. 2007; Fonseca et al. 2012; Reis et al. 2018)). Primers were developed for one breakpoint and its corresponding ancestral state for each of the eight inversions (File S2) based on the *D. virilis* (Clark et al. 2007), *D. americana* (H5, W11, (Fonseca et al. 2013) and SF12, (Reis et al. 2018)), and *D. novamexicana* (15010-1031.00, (Reis et al. 2018)) genome sequences to avoid polymorphism at the primer binding sites. Genomic DNA was extracted from pools of 20 females for each strain using a standard phenol:chloroform protocol, and the concentration was normalized based on concentration measurements using Nanodrop® prior to PCR amplification (File S2). The amplification products were visualized on a UV transilluminator after electrophoresis using TAE buffer in 2% agarose gels stained with a 1:10 dilution of SERVA® stain. Associations between the presence of inversions and eye size (after accounting for body size as described above) were tested using Wilcoxon rank-sum test followed by Holm correction for multiple testing. Associations between mean head shape variation and the presence of inversions were tested using a parametric T-square test on the group mean shapes followed by 10,000 permutations (leave-one-out cross-validation) as implemented in the “Discriminant Function Analysis” option in MorphoJ.

### Parental strains selection and dominance relationships

*D. americana* (SF12) and *D. novamexicana* (15010-1031.00) strains were selected as representatives of both species, because they had the largest differences in eye size (see Results), they show the most divergent karyotypes regarding chromosomal inversions and their genomes are available (Reis et al. 2018). We established several crosses with 10 males and 10 females for each of both parental strains, and 10 *D. novamexicana* males and 10 *D. americana* females to obtain F1 hybrids. Since, *D. americana* females and males take at least four to six days to reach full maturity (Pitnick et al. 1995), we transferred the flies into new vials after seven days. The flies were then allowed to lay eggs for 24h only, to avoid crowding effects on adult size before dissection. Newly eclosed flies were sexed and collected into new vials and were kept under the same conditions described above. Next, 100 females of each parental strain, as well as 100 F1 females were dissected between 4 and 7 days after eclosion, as described above. We used females only to avoid potential confounding effects caused by sex differences (e.g (Siomava et al. 2016)) and to avoid hemizygosity for the *X* chromosome. Pictures were taken using a stereomicroscope Nikon ZMS 1500 H with a magnification of 40x for wings and 50x for the other structures. We also took a picture of a ruler to be able to convert the measurements to μm or μm^2^. The resulting JPG files were saved with a resolution of 1600×1200 pixels. All pictures were treated using ImageJ (Schneider et al. 2012) as mentioned before. After phenotyping, six individuals were excluded from the analysis because they showed highly damaged wings (File S4). We used Kruskal-Wallis test followed by Dunn’s post-hoc test with Holm correction to determine which comparisons were significantly different between strains and hybrids. The scaling relationships between parental strains, as well as between *D. novamexicana* and F1 hybrids were evaluated using the regression of log-transformed eye area (non-corrected for body size) on log-transformed GMsqTW. The slope of the resulting curves is the allometric coefficient (Huxley 1924; Huxley and Teissier 1936). We tested for the significance of differences in allometric coefficients using a linear model with interaction terms.

The geometric morphometric analysis of head shape including removal of artificial pitch and yaw was done as described above (geometric morphometrics section).

For ommatidia counting, the heads of 10 females for each parental strain and hybrids were dissected 4-7 days after eclosion and cut in half longitudinally with a razor blade. One of the eyes of every individual was mounted on sticky tape facing upwards. Serial stack pictures (N=25) were taken using a microscope Zeiss Axioplan 2 with external light sources from the sides and 160X magnification to capture the reflection of every ommatidia (Fig. S1A-B). We also took a picture of a ruler to be able to convert the measurements from pixels to μm or μm^2^. Images were saved with 1360 × 1036 resolution. Stack projection with maximum intensity was achieved using ImageJ (Schneider et al. 2012). The area of the eye was outlined and measured. The images were transformed into 8-bit (gray scale) and the area outside the selected region was cleared. Next, we used the Fast_Morphology.jar plug-in with the following settings: morphological filters, white Tophat – octagon – radius = 2. The images were inverted and the ommatidia numbers were estimated using the ITCN_1_6.jar plug-in with the following settings: width=7px; Minimum distance = 10, threshold=2.0 and detect dark peaks. To estimate the average ommatidia size, the eye area was divided by the number of ommatidia.

### Genotype-phenotype association study using a backcross approach

To determine the effect of the major chromosomes on size variation, we established backcrosses between F1 females (progeny of crosses between *D. novamexicana* males and *D. americana* females) and *D. novamexicana* males. A total of 570 females were dissected and phenotyped for eye, face and wing areas, as well as for tibiae lengths as described above. After phenotyping, 11 females were excluded from the analysis because they showed highly damaged wings (File S4). The remaining 559 females were genotyped using the molecular markers A6 (Muller A, *X* chromosome), B3 (Muller B, *4*^*th*^ chromosome), C3 and C5 (Muller C, *5*^*th*^ chromosome), D7 (Muller D, *3*^*rd*^ chromosome) and E7 (Muller E, *2*^*nd*^ chromosome) (see (Reis et al. 2014) and File S5 for more details). PCR reactions were done using Phire Plant Direct PCR kit^®^ (Thermo Scientific^®^) and gDNA from wings according to the manufacturer’s instructions. We found some unspecific amplification with molecular markers A6 and B3. Based on the recently published *D. americana* (SF12) and *D. americana* (15010-1031.00) genomes (Reis et al. 2018), we were able to slightly modify these primers to account for polymorphisms and improve PCR amplification (File S4). No recombinants were found between molecular markers C3 and C5 on the *5*^*th*^ chromosome (Muller C). The *2*^*nd*^ and *3*^*rd*^ chromosomes (Muller E and D, respectively) are fused in *D. americana*. These are, thus, transmitted as a single chromosome and we found only four recombinants between the molecular markers D7 and E7 (File S4). We excluded the four recombinants and also three individuals that showed the amplification product of *D. americana* only for marker A6 or E7 (File S4). This cleaned data set of 552 females was used to evaluate the effect of each chromosome in eye and wing areas, as well as in GMsqT with the Wilcoxon-rank test followed by Holm correction for multiple testing. Since we observed a strong effect of the *5*^*th*^ chromosome (Muller C) on the size of all analyzed organs (see Results), we decided to use the residuals of the multiple linear regression of eye size on tibiae and wing sizes to account for body size variation. With this conservative approach we removed all the variation in eye size that could be explained by variation in the other structures. We have summed the grand mean of eye size to the residuals to get normalized eye size. Since, normalized eye size is normally distributed (Shapiro-Wilk test, p>0.05), we used linear models with each chromosome as fixed effect to test for significant associations and to estimate the amount of variation explained. We have further included interaction terms between different chromosomes to evaluate epistasis.

The geometric morphometric analysis of head shape was mainly done as described above. In this analysis, pitch and yaw were partly associated with variation along PC1 and PC2, respectively. The error was removed prior to the analysis using the method described before. The impact of the different chromosomes on mean shape variation was evaluated using the “Discriminant Function Analysis” option in MorphoJ. We further used the residuals of the regression of the Procrustes coordinates on centroid size to estimate the impact of the different chromosomes on the non-allometric component of shape. These analyses were also conducted in MorphoJ.

### Genome-wide association study using a pool-seq approach

#### Crosses, sample preparation and sequencing

To identify single nucleotide polymorphisms (SNPs) associated with normalized eye size, we performed a genotype-phenotype association study using F18 individuals resulting from brother sister mating for 17 generations starting from crosses between *D. novamexicana* (15010-1031.00) males and *D. americana* (SF12) females. We phenotyped a total of 157 females, which represented the entire F18 female progeny of two independent vials, for eye and wing areas, as well as tibiae lengths of one of each pair of legs. Wings and legs were randomly dissected from left or right sides. Missing values for individuals ID=125 and 150 showing highly damaged wings were estimated based on the equation of the multiple linear regression between wing area and tibiae length. The residuals of the multiple regression between eye area and both tibiae lengths and wing areas were used to remove the variation in eye area that could be explained by variation in total body size. The grand mean was summed to the residuals and these new values were sorted in ascending order to divide the females into four quartiles (Q1(n=40; 238,913±7,479μm^2^); Q2(n=39; 252,838±2,352μm^2^), Q3(n=38; 263,652±3,777μm^2^) and Q4(n=40; 278,609±9,736μm^2^) (File S4). gDNA for pooled carcasses was extracted for each quartile using a standard phenol:chloroform procedure. DNA quantity and integrity were checked by agarose gel electrophoresis. The good quality samples were used to prepare gDNA libraries for all four pooled samples with TruSeq® Nano DNA Library Prep from Illumina (Catalog#FC-121-9010DOC). The libraries were further used for paired-end sequencing with HiSeq2000 (Illumina) at the Transcriptome Analysis Laboratory (TAL) in Göttingen.

#### Quality checks and analysis

After sequencing using HiSeq2000, a total of 88,783,338; 75,693,540; 82,724,476 and 108,310,437 paired-end reads with 100 bp were obtained for Q1, Q2, Q3 and Q4, respectively. The reads are available at ENAXXXXX. The quality of the reads was assessed with FASTQC v0.11.1 (http://www.bioinformatics.babraham.ac.uk/projects/fastqc/). There was no need to trim or mask positions, since all positions had quality above 20. As mapping reference the *D. americana* (SF12) genome was used after it was reordered based on the hypothetical chromosomes of the ancestral state between *D. virilis*, *D. americana* and *D. novamexicana* (Reis et al. 2018). Read mapping, ambiguously mapped reads and optical duplicates removal, as well as SNP calling and depth of coverage determination were done as described in (Reis et al. 2018). The overall alignment rates using Bowtie2 v2.2.5 (Langmead and Salzberg 2012) with default settings for Q1, Q2, Q3, and Q4 were 77%, 79%, 79% and 81%, respectively. The distributions of coverage obtained with GATK DepthOfCoverage v3.4.46 (Van der Auwera et al. 2013) were close to normal and the average values for Q1, Q2, Q3, and Q4 were 84X, 73X, 81X and 107X, respectively. For all quartiles, more than 94% of the sites showed coverage values above 20X. To avoid bias in frequency estimations, we used a coverage interval which included 68.2% of the total amount of sites around the mean for each quartile ([62-101X], [54-89X], [60-97X] and [84-127X] for Q1, Q2, Q3 and Q4, respectively). The data was treated and analyzed as described in (Reis et al. 2018). Briefly, we used the frequencies values for SNPs identified using both Bowtie2 v2.2.5 (Langmead and Salzberg 2012) and BWA v0.7.12 (Li and Durbin 2009) to determine which ones showed significant differences between the quartiles using Fisher exact test followed by Bonferroni correction.

### Ancestral reconstruction of phenotypic traits

We used the phylogeny of 59 *Drosophila* species obtained by (Keesey et al. 2019) to reconstruct the ancestral state for body size, eye surface, as well as the ratio between eye and head width using a Maximum Likelihood method as implemented in the function fastAnc in the R package Phytools (v. 0.4.98, http://www.phytools.org/eqg2015/asr.html, (Revell 2012)). The details about the strains and the phenotypes can be found in (Keesey et al. 2019).

We also used Phytools to reconstruct the ancestral state of different traits for the species of the *virilis* phylad. The traits considered were the following: GMsqTW as a proxy for body size, eye area, the ratio between eye and head area, and eye area after accounting for body size (residuals of the linear regression between eye area and GMsqTW plus the grand mean of eye area). To obtain phylogenies representative of the *virilis* phylad, we started by downloading all the *D. virilis* coding sequences (CDS) available at FlyBase (ftp://ftp.flybase.net/genomes/Drosophila_virilis/current/fasta/). We further used SEDA (López-Fernández et al. 2019) to retrieve only those CDS of genes located on scaffolds anchored to chromosomes (Muller E: scaffolds 12,822; 13,047; 12,855 and 12,954; Muller B: scaffolds 13,246; 12,963 and 12,723 and Muller C: scaffolds 12,823; 10,324; 12,875 and 13,324). When more than one isoform was available for the same gene only the longest one was retrieved. Next, we used Splign-Compart (as implemented in BDBM; (Vázquez et al. 2019), and the *D. virilis* CDS obtained above as references to annotate CDS in *D. americana* (H5 and W11 (Fonseca et al. 2013); SF12, Northern, Central and Southern populations (Reis et al. 2018)), as well as in *D. novamexicana* (15010-1031.00) contigs (Reis et al. 2018). To obtain the CDS for the *D. americana* populations, we used Coral 1.4 (Salmela and Schroder 2011) with default parameters to reconstruct the gene sequences showing the major frequent variant at polymorphic sites in the pool-seq reads of each population (Reis et al. 2018), prior to contig assembly using Abyss 2.0 (Jackman et al. 2017) with K=25 and default parameters.

For each genome, we used SEDA to filter the datasets for complete CDS (those with annotated start and stop codons) and obtain one file per gene with the orthologous CDS. Sequences were aligned using Clustal Omega (Sievers et al. 2011) and concatenated, resulting in alignments with 372,546 bp (379 genes), 335,931 bp (379 genes) and 298,233 (334 genes) for the *2*^*nd*^, *4*^*th*^ and *5*^*th*^ chromosomes (Muller E, B, and C), respectively. FASTA files were converted to NEXUS format using ALTER (Glez-Peña et al. 2010). The phylogenies were obtained with MrBayes (Ronquist et al. 2012) using the GTR model of sequence evolution allowing for among-site rate variation and a proportion of invariable sites. Third codon positions were allowed to have a different gamma distribution shape parameter than those for first and second codon positions. Two independent runs of 1,000,000 generations with four chains each (one cold and three heated chains) were used. Trees were sampled every 100th generation and the first 2500 samples were discarded (burn-in) (Fig. S11 A-C). The Docker images used for running the above software applications are available at the pegi3s Bioinformatics Docker Images Project (https://pegi3s.github.io/dockerfiles/). Since we have no phenotypic data for *D. americana* H5, W11, and the Central population, their branches were manually removed from the phylogenies. The distances between nodes were accounted for to re-estimate the new branch lengths when applicable (Fig. S11 D-F). We kept the Northern population branch as a phylogenetic proxy for O43 and O53 phenotypes and we decided to remove the Southern population branch because we have genomic data for SF12. The phylogenies were rooted by the *D. virilis* branch prior to ancestral state reconstruction of the size phenotypes using Phytools as described above and ancestral state reconstruction of shape using squared-changed Parsimony (Maddison 1991) as implemented in the option “Map Onto Phylogeny” in MorphoJ.

### Intersection with previous results and SNP annotation

To identify variants associated with normalized eye size and linked with chromosomal inversions, we intersected the SNPs obtained for the F18 pool-seq described above with those obtained in a previously published pool-seq of *D. americana* populations (Reis et al. 2018) (Fig. S8). We started by re-mapping the raw reads obtained for the genome sequencing of *D. americana* (SF12) and *D. novamexicana* (15010-1031.00) against the reference *D. americana* (SF12) genome. Read quality was assessed with FastQC v0.11.1 (http://www.bioinformatics.babraham.ac.uk/projects/fastqc/), and every position showing a quality score under 20 was masked using fastq_masker implemented in the FASTX Tool kit v.0.0.13 (http://hannonlab.cshl.edu/fastx_toolkit/index.html). Read mapping, alignment filtering and SNP calling was done as described above. The mean coverage and standard deviation (s.d.) were determined for both samples and all SNPs showing lower or higher frequency than mean ± 3 s.d. were considered either as errors or originating from highly repetitive regions and were discarded. Next, every SNP showing coverage higher than zero for the reference variant in *D. novamexicana* (15010-1031.00) and the alternative one in *D. americana* (SF12) was excluded. These were variants that might be shared between the two strains, and they would cause biased frequencies. Then, these tables were intersected with those obtained for Q1 and Q4 of the F18 pool-seq described above. Fisher exact test followed by Bonferroni correction was used to determine which SNPs show significant frequency differences between Q1 and Q4. We kept those SNPs which show the reference allele at higher frequencies for Q4. Since, *D. virilis* and *D. novamexicana* show small eye size (see Results), we considered that the variants causing this phenotype would be fixed in those species. The alternative variant should be fixed in *D. americana* (SF12) and at higher frequencies in Q4 and southern populations.

To annotate the SNPs to genes, we started by aligning the *D. americana* (SF12) reference genome to the *D. virilis* genome using Mauve v2.4.0 (Rissman et al. 2009). The raw SNPs coordinates were extracted and were converted according to the position and orientation of the different *D. virilis* scaffolds to match the coordinates provided on the gtf file (ftp://ftp.flybase.net/genomes/Drosophila_virilis/dvir_r1.07_FB2018_05/gtf/). The gtf file was converted to genePred format using gtfToGenePred tools (http://hgdownload.soe.ucsc.edu/admin/exe/linux.x86_64/). A file containing all the transcripts of *D. virilis* was generated using the genePred file and the FASTA file (dvir-all-predicted-r1.07.fasta.gz) with all predicted annotations available for *D. virilis* (ftp://ftp.flybase.net/genomes/Drosophila_virilis/dvir_r1.07_FB2018_05/fasta/) using the script retrieve_seq_from_fasta.pl provided in ANNOVAR v. (Mon, 16 Apr 2018), (http://www.openbioinformatics.org/annovar/annovar_download_form.php, (Wang et al. 2010). The SNPs were then annotated using the script ./table_annovar.pl provided in ANNOVAR.

To determine whether some of the identified SNPs were located in genes associated with eye development, we intersected the list of genes with those described as being involved in “eye development” (GO: 001654; http://flybase.org FB2018_03). We observed that the *D. virilis* orthologs of three genes (*PQBP1*, *CkIIalpha*, and *hyd*) are located on scaffold 12,958. Since this scaffold is not anchored to any Muller element in *D. virilis*, we excluded those genes from the analysis. We also decided to include the manually curated genes *ewg* and *pnr* which have been reported to be involved in eye development (FlyOde; http://flyode.boun.edu.tr/; (Koestler et al. 2015), but are not present in the GO term: “eye development”.

To identify those regions that can explain both intra- and inter-specific variation in normalized eye size, we have further filtered the SNPs which showed significant frequency differences between northern and southern *D. americana* populations after Bonferroni correction. Gene enrichment was tested using Chi-square test with Yates correction. To obtain an estimation of the total number of genes in *D. melanogaster*, we obtained the list of genes in the Geneontology Panther Classification system (http://pantherdb.org/). This list was used to retrieve the chromosomal location in Flybase (http://flybase.org/batchdownload). Only those genes located on Muller A, B, C, D, E and F were considered (File S3).

### Data availability and statistical analysis

All pictures can be found at DRYAD/FIGSHARE (doi, tba), and raw measurements in File S4. All statistical analyses presented in this work were done using R (R Core Team 2018) and the R package Rcmdr (Fox 2005; Fox 2017; Fox and Bouchet-Valat 2018). The plots were prepared using ggplot2 (Wickham 2016) and Microsoft Office. All custom scripts and analysis pipelines are available on the DRYAD/FIGSHARE (doi, tba) repository.

## Supporting information

Descriptive Stats

Primers_Genotyping

SPNs_Intersection_EyeDev Genes

Raw Measurements

Primers_Backcross

## Acknowledgements and Funding

NP, MR, BH and GW were funded by the Deutsche Forschungsgesellschaft (DFG, Grant Number: PO 19 1648/3-1) to NP. MR was funded by the Volkswagen Foundation (Support for Europe, Grant Number: 85983-1) to NP and JV. Many thanks to the Deep-Sequencing Core Facility of the Universitätsmedizin Göttingen (UMG) for next generation sequencing.

## Authors contribution

**MR:** Conceptualization, planning and execution of the experiments (fly husbandry, handling and crossing, dissections, mounting and pictures capturing, gDNA extraction for PoolSeq); data analysis: size (script writing for statistical analysis using R), shape (handle tps files, create sliding LM, R scripting to remove error, and statistical analysis in MorphoJ), ancestral reconstruction (phytools and MorphoJ); Poolseq analysis (script writing for read quality checking, and mapping, applying the scripts written by CR and HNT for SNP calling and statistical analysis, script writing for table intersection to obtain relevant SNPs); Writing – original draft, Writing – review and editing

**GW:** development of molecular markers for chromosomal inversions, DNA extraction and genotyping

**JC:** support with the geometric morphometrics analysis, reviewing and editing the manuscript

**RL:** manual measurements of heads, eyes, tibiae and wings

**BH:** placing LM for geometric morphometrics analysis

**CR:** script writing for variant calling of the Poolseq GWAS data with GATK

**NTH:** script writing for statistical analysis (Fisher exact test) of the Poolseq GWAS data and data visualization (Manhattan plots)

**CPV:** Conceptualization, review and editing manuscript

**JV:** Conceptualization, Funding acquisition, Phylogeny reconstruction of the virilis phylad, review and editing of the manuscript

**NP:** Conceptualization, Funding acquisition, Project administration, Resources, Supervision, Visualization, Writing – original draft, Writing – review and editing.

## List of supplementary figures

**Fig. S1. Differences in eye size, ommatidia number, and ommatidia size between parental strains and their interspecific hybrid.**

**Fig. S2. Variation in organ size in the genotype-phenotype associations using the backcross approach.**

**Fig. S3. Variation in eye size and head shape in the genotype-phenotype associations using the backcross approach.**

**Fig. S4. Normalized eye size variation for the 16 genotypic classes present in the backcross between F1 hybrid females and** *D. novamexicana* **males.**

**Fig. S5. Manhattan plots between adjacent quartiles of the pool-seq experiment involving F18 females.**

**Fig. S6. Phylogeny and ancestral reconstruction of the strains used in this study.**

**Fig. S7. Comparison between the ancestral reconstruction of phenotypic traits across the** *Drosophila* **genus and the strains used in this study.**

**Fig. S8. Flowchart summarizing the procedure used to obtain the list of relevant SNPs.**

**Fig. S9. Schematic representation of the procedure used for phenotyping.**

**Fig. S10. Sequential removal of error associated with head tilting.**

**Fig. S11. Phylogenies of species of the** *virilis* **phylad.**

## List of files

**File S1. Descriptive statistics for all datasets.**

**File S2. List of primers used as molecular markers for chromosomal inversions and genotyping results.**

**File S3. SNP tables after intersecting the datasets obtained for the GWAS and** *D. americana* **populations and candidate genes for eye development.**

**File S4. Raw measurements.**

**File S5. List of primers used as indel markers for the different chromosomes and genotyping of the progeny of the backcross between hybrid females and** *D. novamexicana* **males.**

