## Supplementary Figures for "Multiple loci linked to inversions are associated with eye size variation in species of the *Drosophila virilis* phylad"

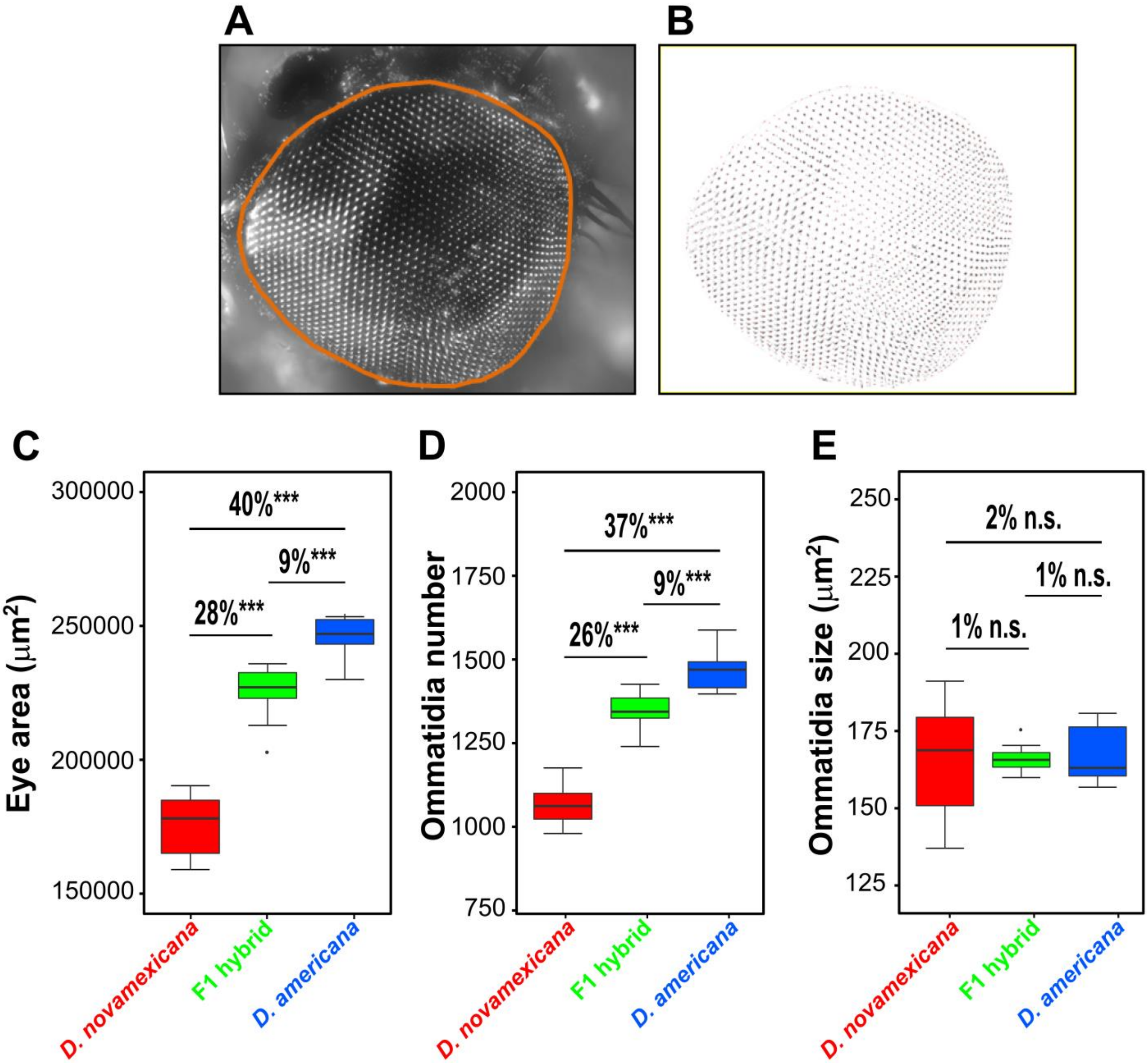


**Fig. S1. Differences in eye size, ommatidia number, and ommatidia size between parental strains and their interspecific hybrid. A.** The measured eye area is depicted in orange. **B.** Transformed pictured used to count ommatidia. **C.** Variation in eye area among parents (10 females each) and hybrids (10 females). **D.** Variation in ommatidia number for the same individuals. **E.** Average ommatidia size obtained as the ratio between eye area and ommatidia number.

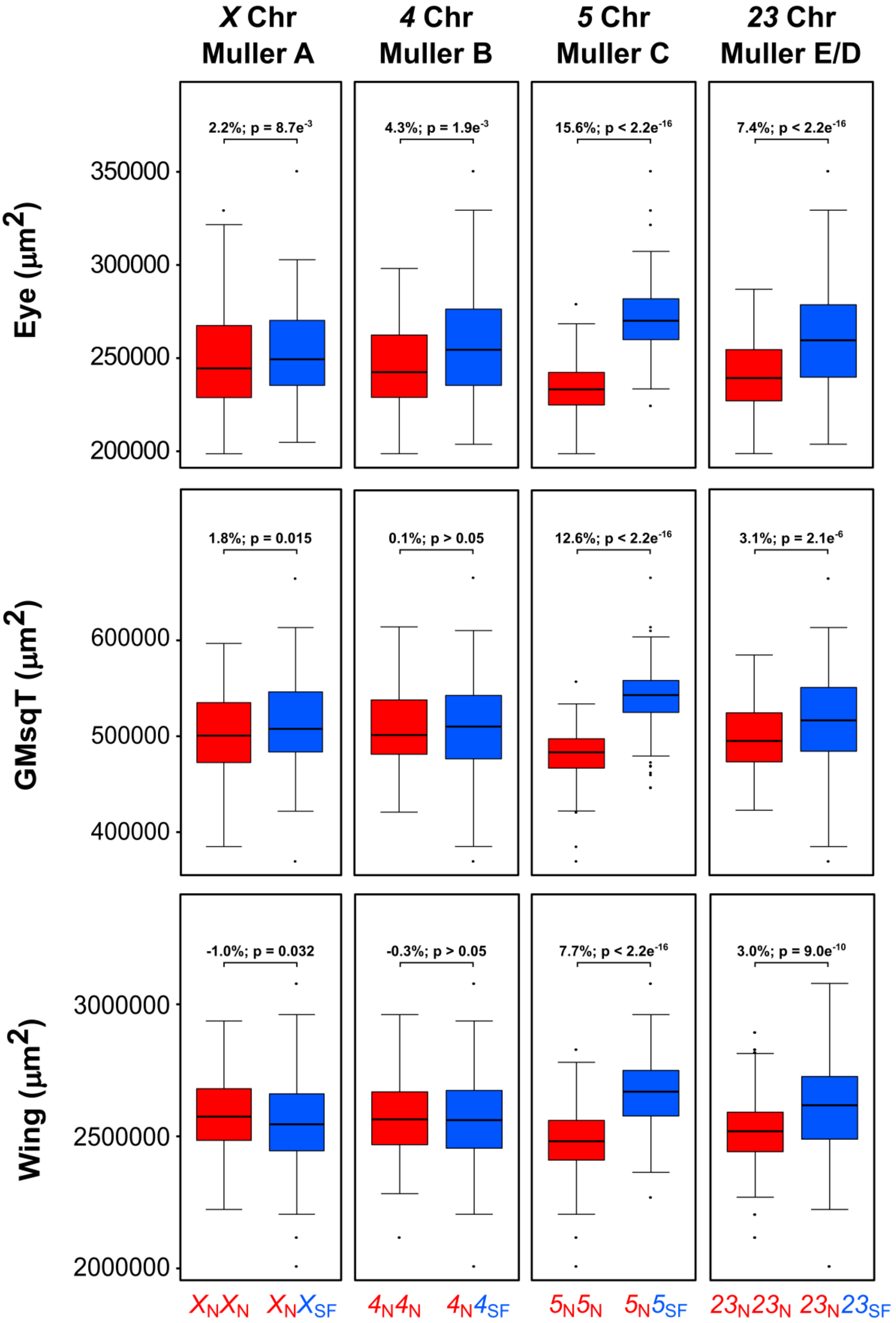


**Fig. S2. Variation in organ size in the genotype-phenotype associations using the backcross approach.** Distributions of eye area, tibiae size (geometric mean of squared tibiae lengths (GMsqT)), and wing area for individuals homozygous for a given *D. novamexicana* chromosome and heterozygous *D. novamexicana*/*D. americana* for the respective chromosome. For each comparison the percentage of difference and significance values after Wilcoxon-rank test are provided.

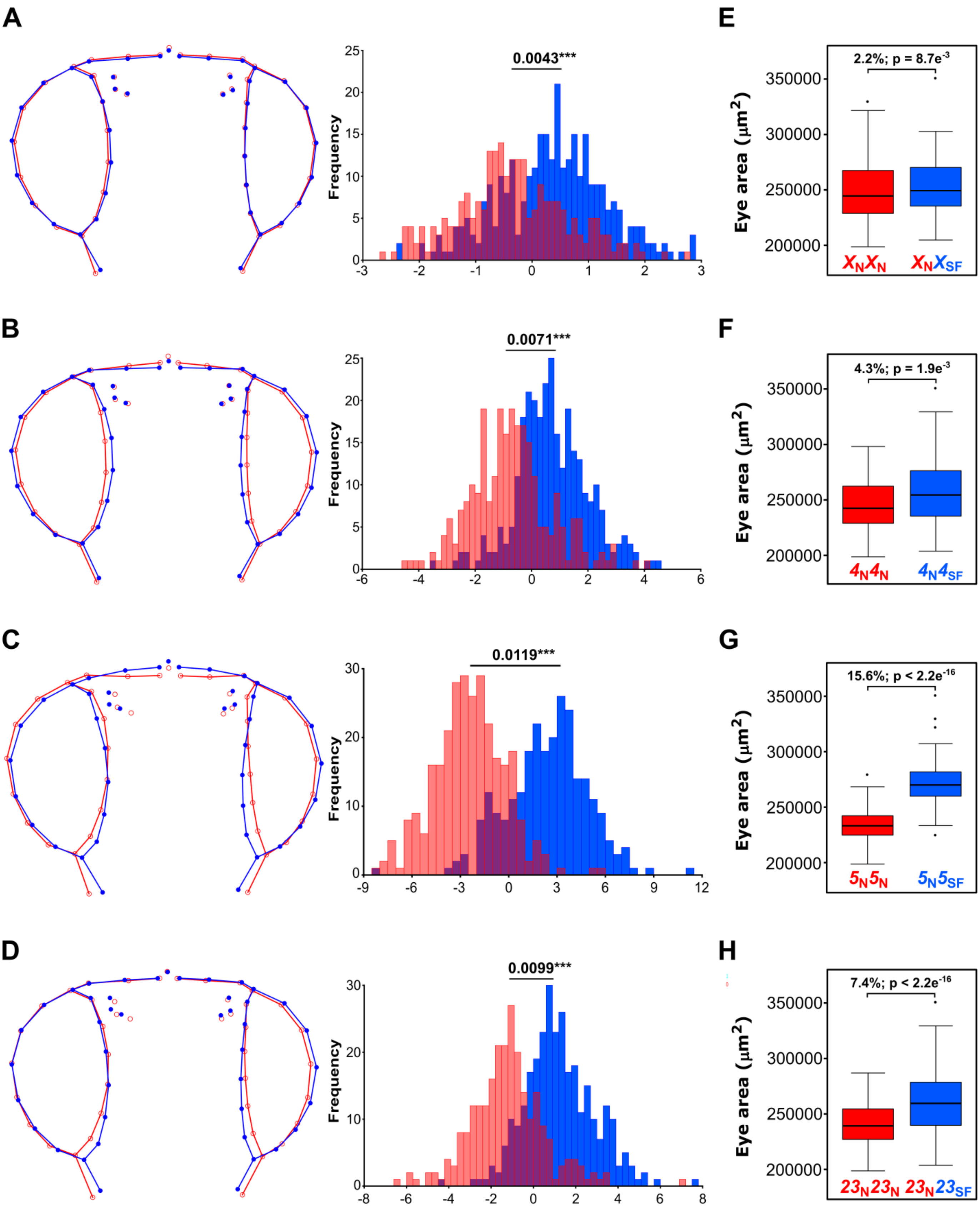

A
B
C
D
E
F
G
H
0.0043***
0.0071***
0.0119***
0.0099***
Frequency
2.2%; p = 8.7e-3
4.3%; p = 1.9e-3
15.6%; p < 2.2e-16
7.4%; p < 2.2e-16
Eye area (µm2)
XNXN
XNXSF
4N4N
4N4SF
5N5N
5N5SF
23N23N
23N23SF

**Fig. S3. Variation in eye size and head shape in the genotype-phenotype associations using the backcross approach. A-D.** Variation in mean head shape among the female progeny of the backcross between F1 hybrid females and *D. novamexicana* males (Discriminant function analysis of the procrustes coordinates obtained from the first 19 principal components (90.8% of the total variation); procrustes distances are provided with *** = p<0.0001 after a permutation test with 10,000 iterations). The wireframes depict changes in the mean shape multiplied by a factor of 5 along the axis of Mahalanobis distances (homozygous *D. novamexicana* (red) or heterozygous *D. novamexicana*/*D. americana* (blue)). **E-H** Distributions of eye size for females, progeny of the backcross, which were homozygous for a given *D. novamexicana* chromosome (red) or heterozygous *D. novamexicana*/*D. americana* (blue) for the respective chromosome. Information about the magnitude of change in eye size and the significance values is shown inside the graphs.

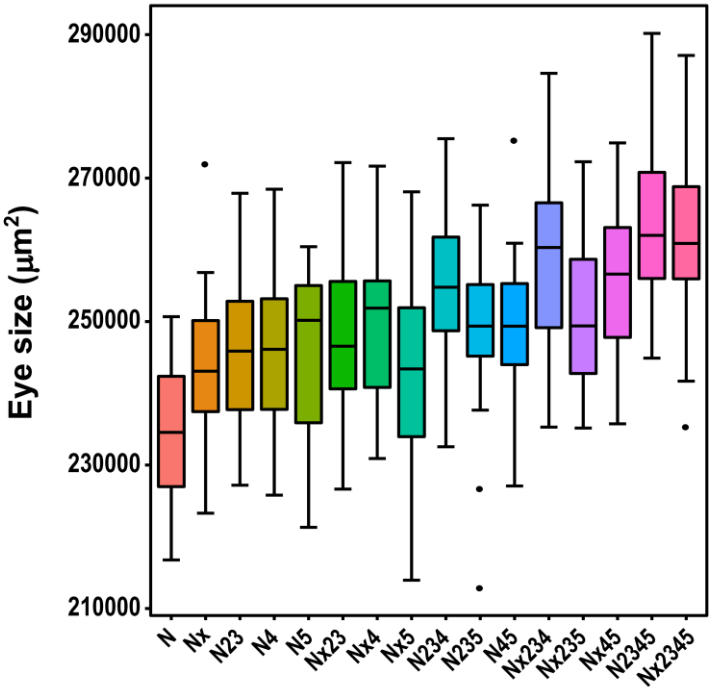


**Fig. S4. Normalized eye size variation for the 16 genotypic classes present in the progeny of backcross between F1 hybrid females and *D. novamexicana* males.** The residuals of the linear regression between eye area and tibiae lengths and wing areas were used to account for variation in body size. The grand mean of eye area was summed to the residuals to get normalized eye size.

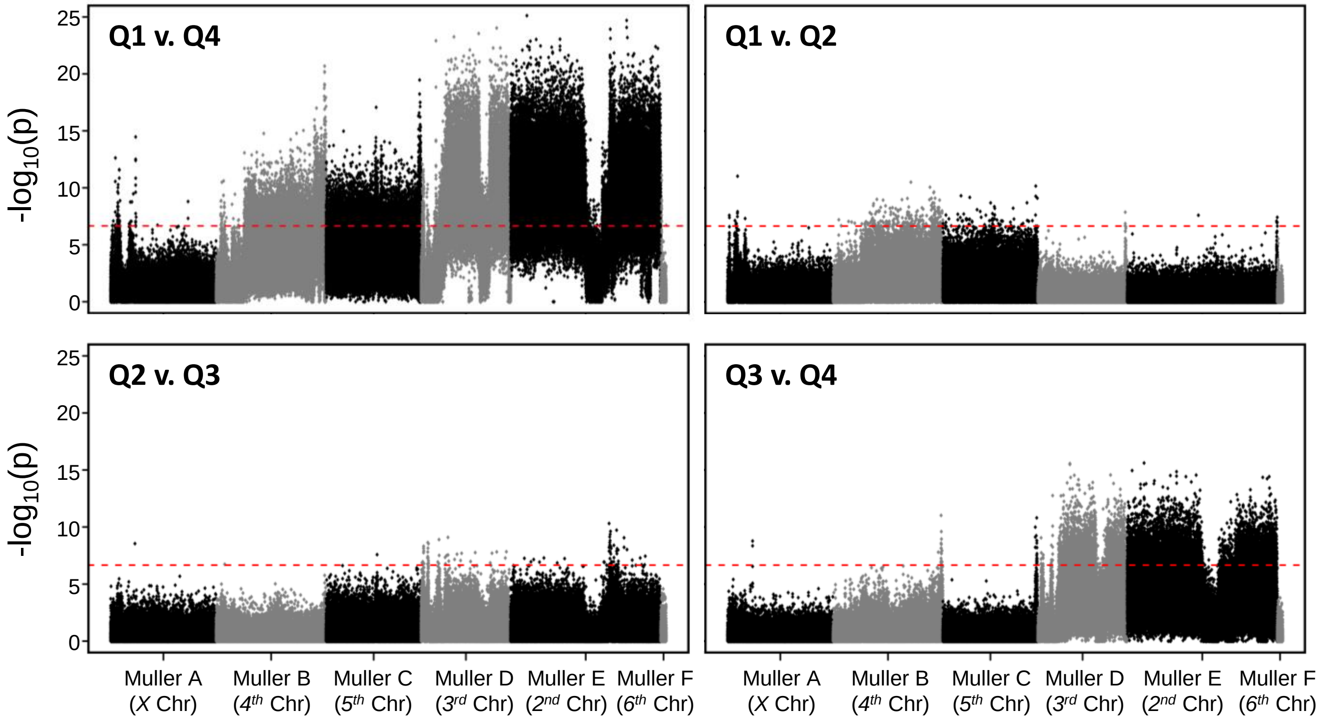


**Fig. S5. Manhattan plots between adjacent quartiles of the pool-seq experiment involving F18 females.** The - $\log_{10}(p)$ values obtained after Fisher exact test were plotted in function of the location in the genome. The chromosomes are identified on the *x* axis and they are oriented always from telomere to centromere. The red lines represent the significant threshold after Bonferroni correction for multiple testing.

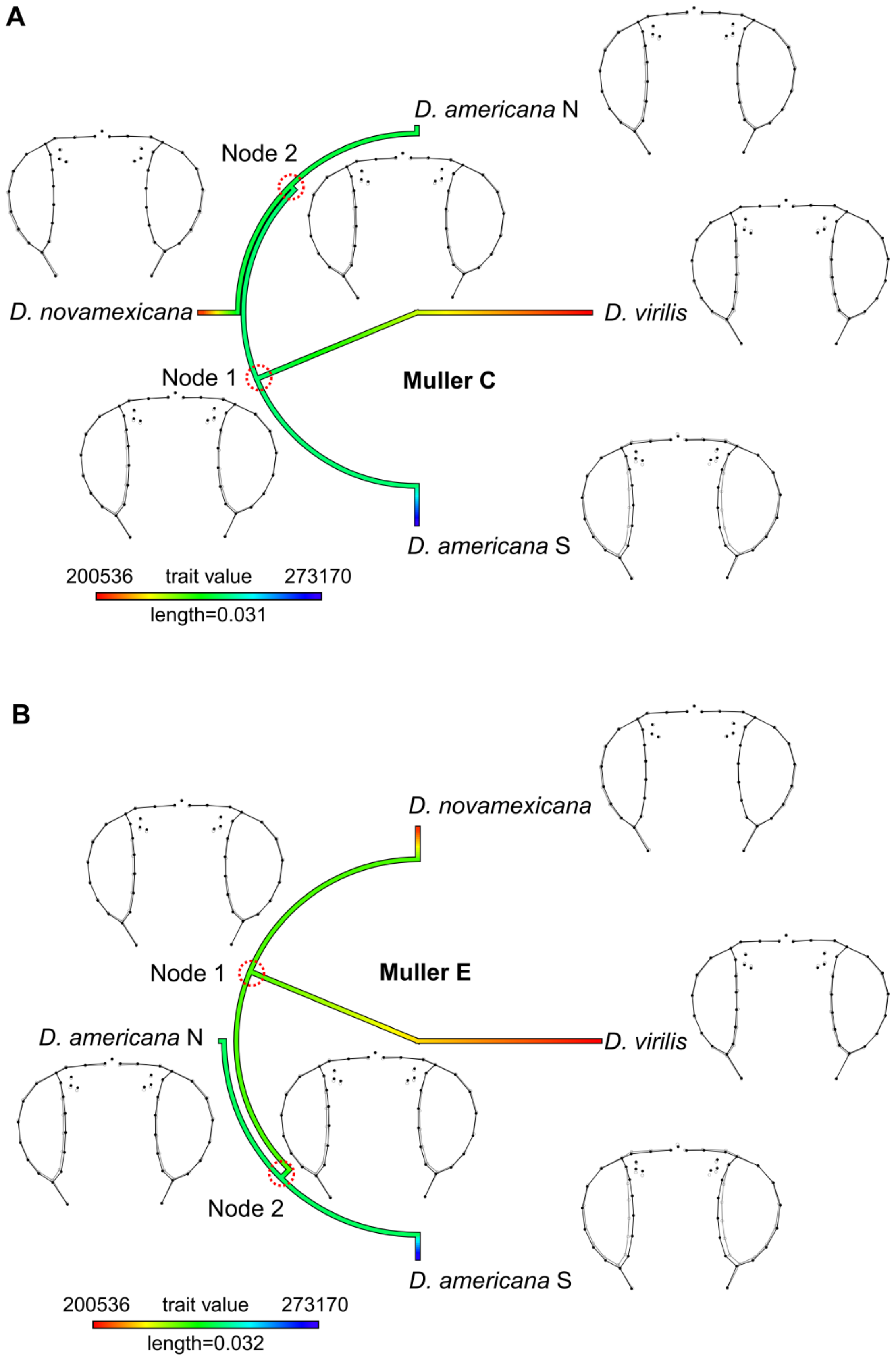


**Fig. S6. Phylogeny and ancestral reconstruction of the strains used in this study. A.** Phylogeny based on genes located on the *5th* chromosome (Muller C). **B.** Phylogeny based on genes located on the *2nd* chromosome (Muller E). The wireframes (black – mean head shape of each species/population, grey – mean head shape of the estimated ancestral) are shown for each species/population.

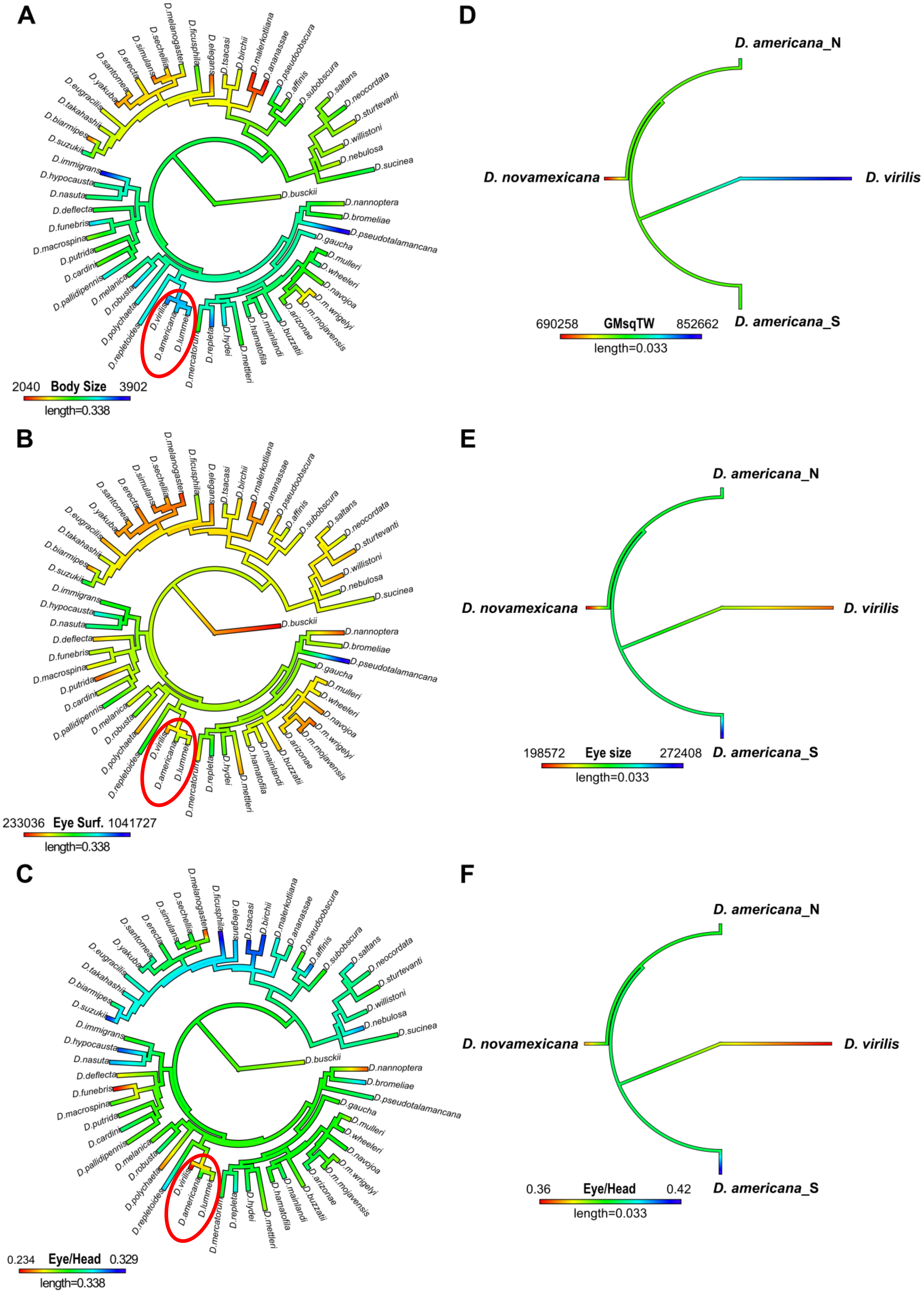
A
D.melanogaster
D.sechellia
D.simulans
D.erecta
D.santomea
D.yakuba
D.eugracilis
D.takahashii
D.biarmipes
D.suzukii
D.ficusphila
D.elegans
D.tsacasi
D.birchii
D.malerkotliana
D.ananassae
D.pseudoobscura
D.affinis
D.subobscura
D.saltans
D.neocordata
D.sturtevanti
D.willistoni
D.nebulosa
D.sucinea
D.busckii
D.immigrans
D.hypocausta
D.nasuta
D.deflecta
D.funebris
D.macrospina
D.putrida
D.cardini
D.pallidipennis
D.melanica
D.robusta
D.polychaeta
D.repletoides
D.virilis
D.americana
D.lummei
D.mercatorum
D.repleta
D.hydei
D.mettleri
D.hamatofila
D.mainlandi
D.buzzatii
D.arizonae
D.m.mojavensis
D.m.wrigelyi
D.navojoa
D.wheeleri
D.mulleri
D.gaucha
D.pseudotalamancana
D.bromeliae
D.nannoptera
2040 Body Size 3902
length=0.338
B
233036 Eye Surf. 1041727
length=0.338
C
0.234 Eye/Head 0.329
length=0.338
D
D. americana_N
D. novamexicana
D. virilis
D. americana_S
690258 GMsqTW 852662
length=0.033
E
D. americana_N
D. novamexicana
D. virilis
D. americana_S
198572 Eye size 272408
length=0.033
F
D. americana_N
D. novamexicana
D. virilis
D. americana_S
0.36 Eye/Head 0.42
length=0.033

**Fig. S7. Comparison between the ancestral reconstruction of phenotypic traits across the *Drosophila* genus and the strains used in this study. A-C.** The phylogeny and phenotypic data for body size (length from thorax to abdomen) (A), eye surface calculated from eye height and width (B) as well as eye length to head length ratios (C) were obtained from Keesey et al. (2019). The red ellipses depict species of the *virilis* group (*D. virilis*, *D. lummei* and *D. americana*). **D-F**. The phylogeny of the *virilis* group based on nucleotide sequences of genes located on the *4$^{th}$* chromosome (Muller B). The phenotypes used for ancestral reconstruction were the following: geometric mean of tibia lengths and wing areas (proxy to body size) (D), eye area measured as the sum of the area of both eyes (E), and the ratios between eye area and head area (F).

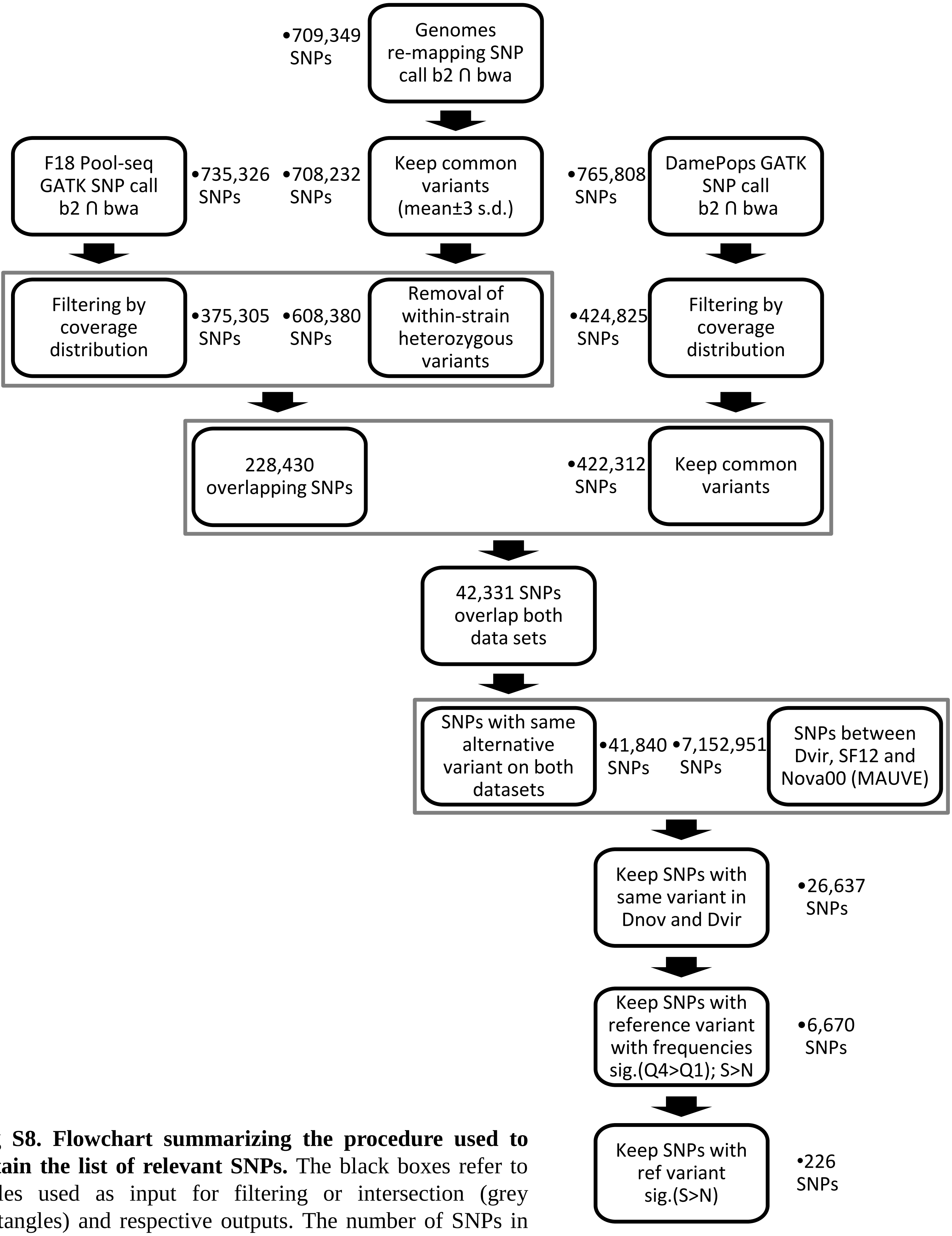


**Fig S8. Flowchart summarizing the procedure used to obtain the list of relevant SNPs.** The black boxes refer to tables used as input for filtering or intersection (grey rectangles) and respective outputs. The number of SNPs in each table is provided next to the boxes. The black arrows depict the different steps (filtering or intersection).

A

B

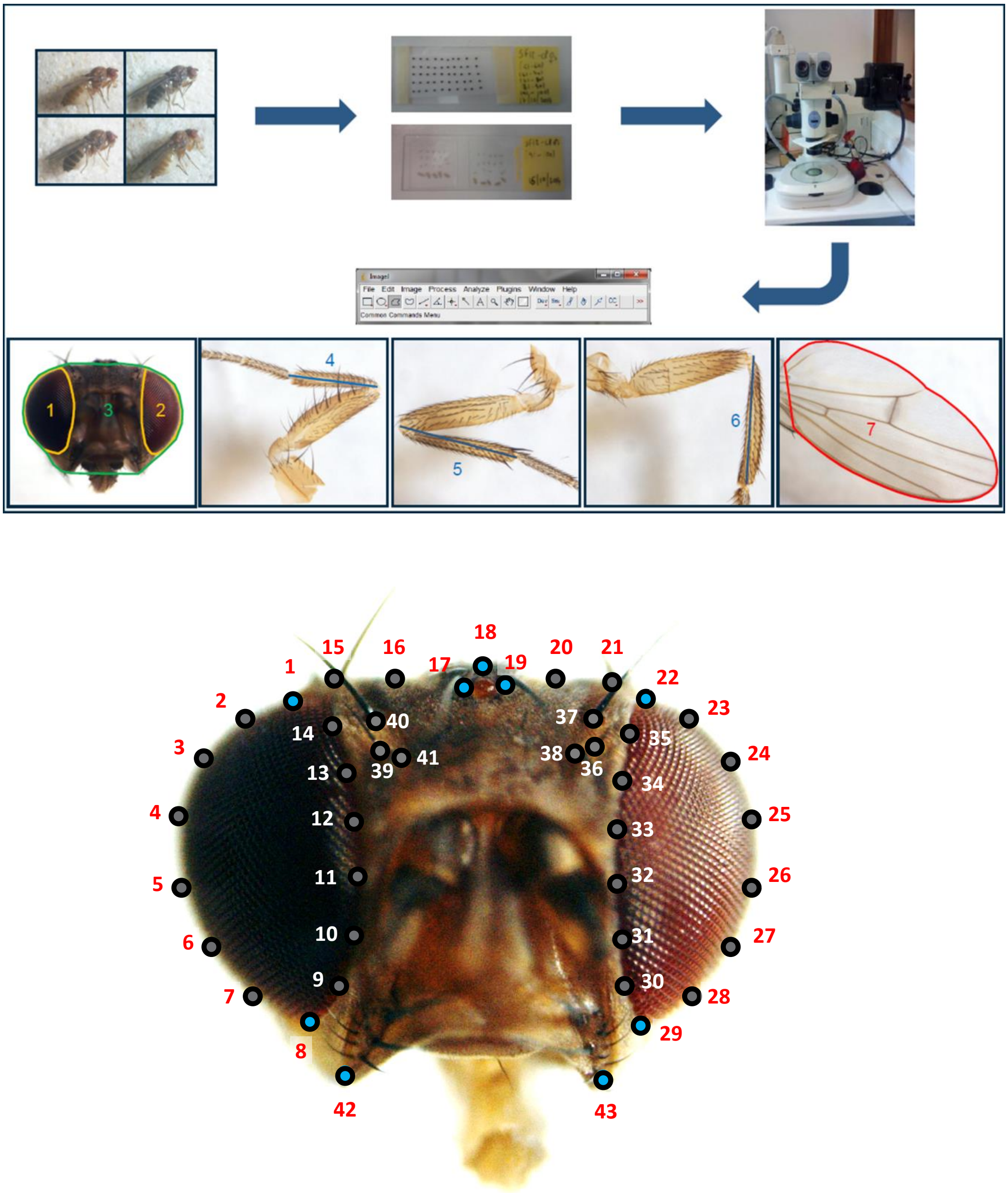


**Fig. S9. Schematic representation of the procedure used for phenotyping. A.** Fly heads were dissected and mounted on sticky tape facing upwards. Tibiae and wings were mounted in Hoyer's medium after dissection. Pictures were taken using a camera attached to a stereomicroscope. Eye area was determined by the area defined by the outlines 1 and 2, while the interstitial head cuticle was determined by subtracting eye area from complete head area [3-(1+2)]. Tibiae lengths were measured as the distances represented by 4, 5, and 6 for tibia 1, 2 and 3, respectively. Wing area was calculated by measuring the area defined by the outline represented by 7. **B.** A total of 11 fixed landmarks (blue) and 32 semi-landmarks (grey) were placed on frontal pictures of heads for geometric morphometrics analysis (see Material and Methods for details).

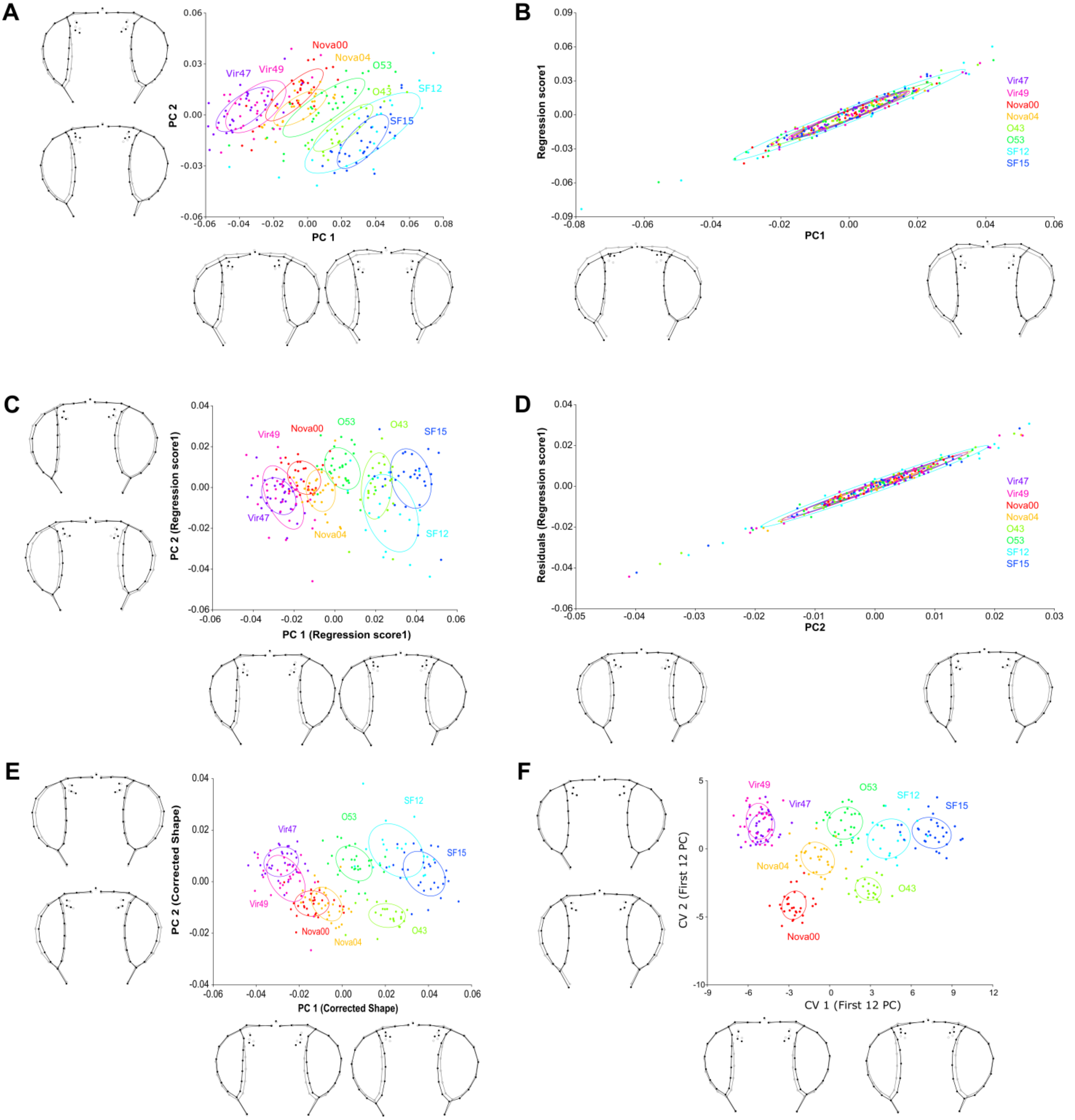

**Fig. S10. Sequential removal of error associated with head tilting. A.** Principal Component Analysis (PCA) of shape of species of the *virilis* group. **B.** Strain-centred regression of shape on PC1 capturing variation associated with roll (up/down). **C.** PCA of the residual variation of shape on PC1 (shape corrected for roll). **D.** Strain-centred regression of shape corrected for roll on PC2 capturing variation associated with yaw (left/right). **E.** PCA of the residual variation of shape corrected for roll on PC2 (shape corrected for yaw). **F.** Canonical Variate Analysis of  shape corrected for roll and yaw. The wireframes depict changes in shape along the two main axes of variation (black - the maximum and minimum values on the axis (Mahalanobis distances); grey – mean shape for each axis). The equal frequency ellipses are given with probability of 0.5.

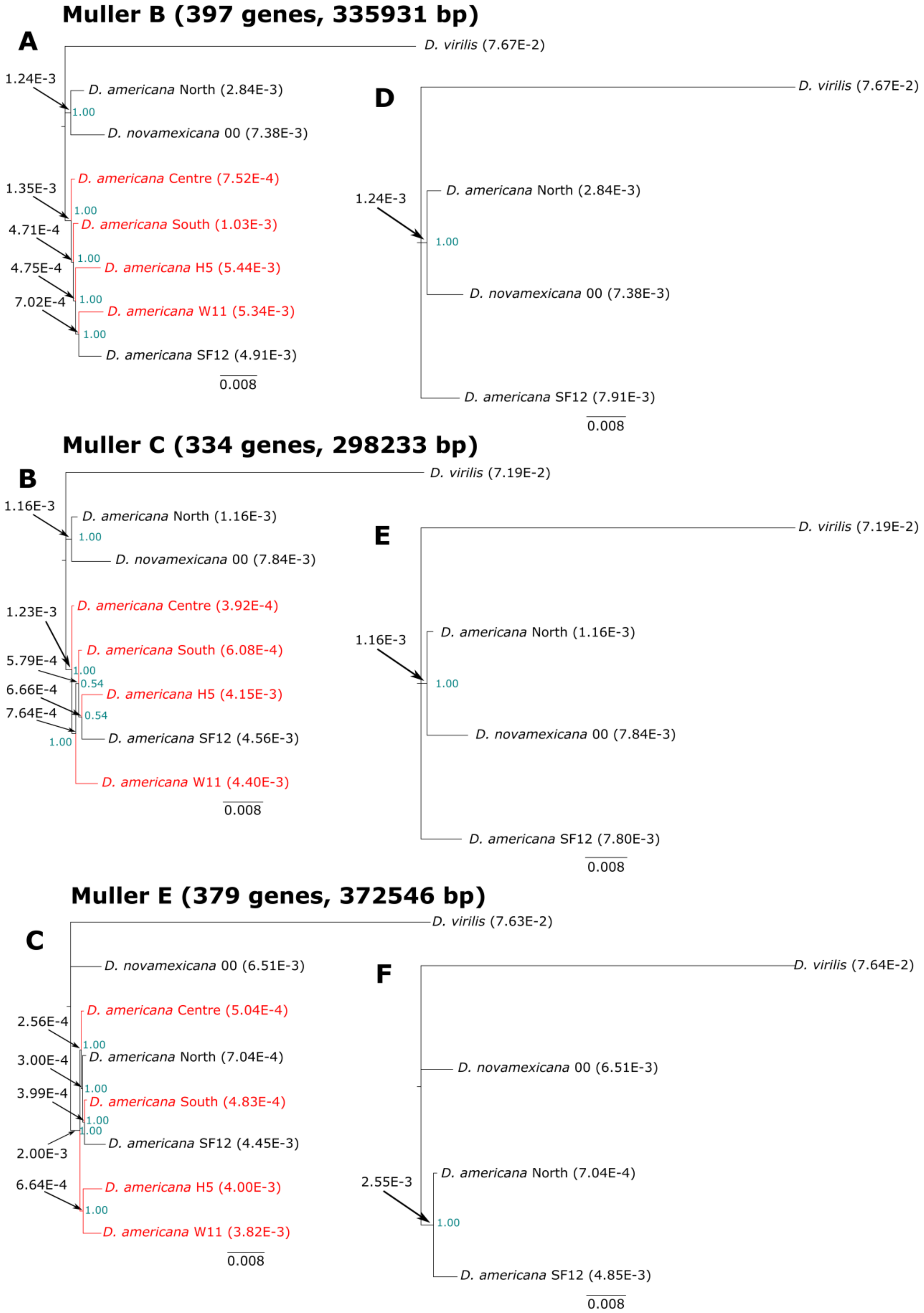


**Fig. S11. Phylogenies of species of the *virilis* phylad. A-C.** Unrooted phylogenies based on genes on the *4th*, (**A**) *5th*, (**B**), and *2nd* (**C**) chromosomes (Muller elements B, C, and E). **D-F.** Edited phylogenies to include only the strains/populations used in this study for the *4th*, (**D**) *5th*, (**E**), and *2nd* (**F**) chromosomes.

### List of additional supplementary files (provided as Excel files)

**File S1. Descriptive statistics for all datasets.**

**File S2. List of primers used as molecular markers for chromosomal inversions and genotyping results.**

**File S3. SNP tables after intersecting the datasets obtained for the GWAS and D. americana populations and candidate genes for eye development.**

**File S4. Raw measurements.**

**File S5. List of primers used as indel markers for the different chromosomes and genotyping of the progeny of the backcross between hybrid females and D. novamexicana males.**
